# Lipid-stabilized ICG Nanoaggregates for the Photodisruption of Vitreous Opacities

**DOI:** 10.1101/2025.09.15.675130

**Authors:** Pouria Ramezani, Jan Félix, Mariana Hugo Silva, Ine Lentacker, Rein Verbeke, Kevin Braeckmans, Stefaan C. De Smedt, Félix Sauvage

**Author notes:** Corresponding address: Ottergemsesteenweg 460, 9000 Ghent, Belgium.

## Abstract

Collagen aggregation in the vitreous is a major cause of vision impairment. Current treatments such as vitrectomy or YAG laser vitreolysis remain limited by invasiveness and safety concerns. In previous work, we introduced a novel approach combining indocyanine green (ICG) with nanosecond laser pulses to achieve photodisruption of collagen aggregates via vapor nanobubbles (VNBs), while using a significantly lower total light dose than that applied in clinical laser vitreolysis. However, despite its clinical approval, free ICG poses a risk of retinal toxicity. In this work, we report the development of ICG nanoaggregates (ICG AGG NPs) stabilized with a minimal amount of a hyaluronic acid (HA)-lipid (DOPE) conjugate designed to limit retinal penetration of ICG while preserving efficient VNB generation and collagen aggregate disruption. We demonstrate that supramolecular aggregation is a key requirement for efficient VNB generation, whereas encapsulation of ICG in conventional liposomes impairs this process. Using a newly established in vitro model for quantifying collagen disruption, we show that ICG AGG NPs significantly enhance photodisruption compared to free ICG. Furthermore, cell toxicity assays on retinal pigment epithelium (RPE) and Müller cells indicate that ICG AGG NPs maintain an acceptable safety profile at therapeutic concentrations. These findings represent the first successful demonstration of dye-loaded nanoparticles enabling efficient VNB-mediated photodisruption of vitreous opacities and highlight the promise of ICG AGG NPs as a safer and more effective alternative to free ICG for floater treatment.

## Introduction

The vitreous humour is the transparent gel which fills the inside of the eye and mainly consists of water, collagen and hyaluronic acid (HA). The collagen of the vitreous maintains its integrity via thin fibrils forming a stable network. These collagen fibrils are believed to remain spaced apart due to electrostatic repulsions facilitated by anionic hyaluronan molecules bound to the collagen fibrils [1, 2]. However, conditions such as myopia and aging can lead to a progressive dissociation of hyaluronan from this network, disrupting the spatial organization of the fibrils. Protein aggregation can then occur, contributing to the formation of collagen opacities in the vitreous body of the eye—a condition which is often overlooked [3] (Figure 1A). These vitreous opacities, composed predominantly of type II collagen, aggregate over time, resulting in patients experiencing ‘floaters’. This phenomenon can cause vision disturbances and reduced quality of life due to the persistent perception of floating shadows or silhouettes in front of the retina [1, 4].

**Figure 1.**
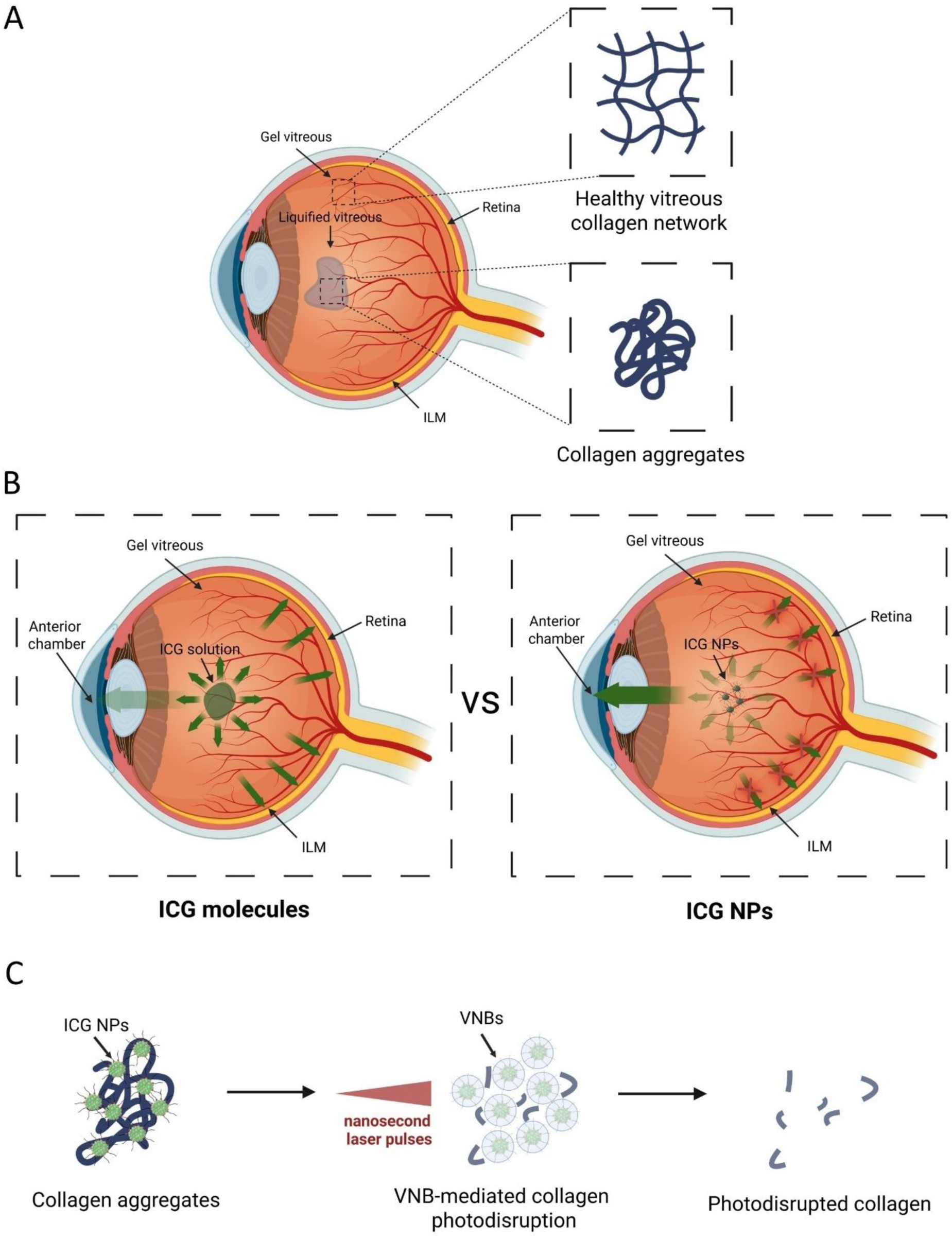
**A)** In the healthy human vitreous, stable collagen strands form a network. In contrast, areas of liquefied vitreous contain aggregated collagen, which may contribute to visual disturbances if aggregates become large enough. **B)** Illustration of the potential advantages of ICG NPs compared to free ICG in terms of retinal safety. Free ICG is expected to diffuse rapidly from the injection site and penetrate through the retina, which may increase the risk of retinal toxicity. In contrast, ICG NPs, due to their larger size, are expected to diffuse more slowly in the vitreous, favoring elimination through the anterior chamber. Additionally, the inner limiting membrane (ILM) could act as a barrier that limits NP penetration, also improving retinal safety. **C)** ICG NPs are irradiated with nanosecond laser pulses to trigger VNB-mediated photodisruption of collagen aggregates. Image created with BioRender.com.

Current treatment strategies include pars plana vitrectomy, in which the vitreous opacities are removed through the removal of the vitreous and YAG laser vitreolysis in which intense (nanosecond) laser pulses are employed to destroy the vitreous opacities [5, 6]. However, both strategies have shortcomings. Vitrectomy is an invasive procedure, often associated with cataract formation [7] while YAG laser vitreolysis requires the use of high-energy laser pulses, which may harm the retina [8, 9], the crystalline lens and the lens capsule [9–11]. As a result, this technique is typically limited to treating vitreous opacities located centrally within the vitreous body—sufficiently distant from both the lens and the retina—to minimize the risk of collateral damage. Additionally, some reports suggest the occurrence of open-angle glaucoma due to an increase in intraocular pressure (IOP) induced by the intense energy levels used in YAG laser therapy [9, 12]. Given these drawbacks and the fact that YAG laser vitreolysis is mostly effective on rather larger and highly dense opacities (e.g. Weiss rings) [13] while showing low effectiveness on other types of opacities [14], a safer, yet more effective, approach for floater treatment, specifically for smaller opacities, is required.

Previously, our group introduced a novel strategy for eye floater treatment using photosensitzers (PS) in combination with nanosecond pulsed-laser light [15, 16]. PS are molecules or supramolecular assemblies that strongly absorb laser light. Upon excitation, they can dissipate their excess energy through non-thermal and thermal pathways which enables biomedical theranostic modalities including fluorescence and photoacoustic imaging as well as photodynamic and photothermal therapy—areas that have received significant attention in recent years [17]. More recently, photodisruption has emerged as a novel modality which relies on the generation of photomechanical forces by the PS. In this modality, excitation of the PS with a pulsed-laser generates intense localized heat, causing the surrounding water to evaporate, leading to the generation of vapor nanobubbles (VNBs) [18]. These VNBs expand and collapse, generating mechanical forces which can disrupt nearby protein aggregates, biological barriers or bacterial biofilms [15, 16, 19, 20]. Previously, we have shown that laser-induced VNBs can destroy collagen opacities in the vitreous with a total light dose 10^3^ to 10^6^ times lower compared to ablation with a clinical YAG laser [15, 16]. In these studies, as PS for VNB generation, we used gold nanoparticles (NPs) [15] and molecular indocyanine green (ICG) [16]. However, gold NPs may fragment under laser illumination, potentially causing genotoxicity [21]. ICG, while FDA-approved and widely used as an ocular dye, can exhibit dark- and light-induced retinal toxicity upon penetration in the retina [22–24]. Therefore, it is essential to improve the safety of VNB-mediated floater destruction by designing new types of PS.

Specifically, we considered formulating ICG into NPs to prevent its free diffusion into the retina, thereby improving its safety profile (Figure 1B). In the context of ocular delivery, NP formulations have been explored for decades, and their application for drug or gene delivery continues to attract significant interest. However, effective retinal delivery of NP formulations upon intravitreal (IVT) administration remains challenging. Following IVT injection, NPs must traverse the vitreous body [25], cross the inner limiting membrane (ILM) covering the retina, and penetrate the retinal layers to deliver their cargo efficiently [26]. Besides, IVT-injected NPs are often hindered within the vitreous and ultimately cleared via the anterior chamber [27]. Moreover, due to the presence of the ILM which acts as a size selective barrier, NPs larger than 100 nm typically fail to penetrate the retina [28]. In contrast, small molecules diffuse more rapidly and cross these barriers more easily, leading to their increased distribution into the retinal tissue and clearance through posterior routes [29, 30]. Thus, in this work, we investigate the formulation of ICG into NPs to enhance safety by limiting retinal distribution and minimizing toxicity. To ensure good intravitreal mobility, we aimed to coat the NPs either with hyaluronic acid (HA) or poly(ethylene) glycol (PEG). HA, being negatively-charged, improves diffusional mobility of NPs in the vitreous while maintaining NP affinity to collagen fibers [15, 31].

In this study, we explore the use of NP-formulated ICG functionalized with PEG or HA to achieve optimal VNB-mediated photodisruption of vitreous opacities (Figure 1C). Two NPs designs are investigated: ICG-loaded liposomes (ICG LIPs) and ICG nanoaggregates (ICG AGG NPs). Our goal is to identify which factors are needed for efficient VNB generation and collagen aggregate photodisruption. We also introduce an *in vitro* model to better quantify the extent of photodisruption based on scattering intensity determined by dark field microscopy.

## Materials and methods

### Materials

1,2-dioleoyl-sn-glycero-3-phosphoethanolamine (DOPE), 1,2-dioleoyl-3-trimethylammonium-propane (DOTAP), 1,2-Distearoyl-sn-glycero-3-phosphocholine (DSPC) and 1,2-distearoyl-sn-glycero-3-phosphoethanolamine-N-[poly(ethylene glycol)-2000] (DSPE-PEG2000Da) lipids were purchased from Avanti Polar lipid, USA. Indocyanine green (ICG) and 20kDa hyaluronic acid (HA) were purchased from Sigma Aldrich, USA. Collagen type I (rat tail) and 6-8kDa SpectraPor dialysis membranes were purchased from ThermoFisher Scientific, USA. 2-Chloro-4,6-dimethoxy-1,3,5-triazine (CDMT) and 4-Methylmorpholine (MM) reagents were purchased from TCI, Japan.

### Synthesis of DOPE conjugated to HA

DOPE was conjugated to 20 kDa HA by using CDMT and MM as coupling reagents, similar to a report by Yoo et al. [32]. Firstly, 50 mg HA (0.125 mmol −COO^−^) was dissolved in 3 ml deionized water (DIW) by adding HA to water, slowly under 250 rpm magnetic stirring (Heidolph MR 3001K, Germany). Afterwards, 4.39mg CDMT (0.025 mmol) and 3.79 mg MM (0.0375 mmol) were dissolved in 1 ml acetonitrile and added to the HA solution in water. This mixture was then stirred at room temperature for 2 h so that the carboxylic groups of HA become activated for the coupling reaction. Next, an ethanolic DOPE solution containing 13.02 mg DOPE (0.0175 mmol) in 4 ml ethanol was added to the activated HA solution under 250 rpm stirring, and this reaction was set for 24 h. Finally, the obtained product was dialyzed two times, first by using 2 L of a 1:1 water/ethanol solution using 6-8 kDa regenerated cellulose (RC) membranes for 24 h and then, for 24 h, only in water, to ensure a purified product. The purified product was freeze-dried and stored in −20°C. A 4 mg/ml solution in D_2_O of the final product was also characterized by ^1^H NMR (see method in the supplementary information and Figure S1).

### Preparation and characterization of HA-coated and PEG-coated ICG liposomes

Both PEG-coated ICG liposomes (ICG-PEG LIPs) and HA-coated ICG liposomes (ICG-HA LIPs) were prepared using the thin-film rehydration method as previously described [33] and summarized in Scheme 1. For both HA-coated and PEG-coated LIPs, an ethanolic solution of a certain amount of lipids (Table 1) was added to a round-bottom flask along with 0.66 mg ICG. A rotary evaporator (IKA RV10, Germany) was used to evaporate the organic solvent and form a thin lipid film under the following conditions: 30 minutes at 40 °C, 250 rpm (Heidolph MR 3001K, Germany), and 50 mbar vacuum. The resulting lipid film was rehydrated with either 1 ml of deionized water for ICG-PEG liposomes (ICG-PEG LIPs) or 1 ml of a 2.5 mg/ml DOPE-HA aqueous solution for ICG-HA liposomes (ICG-HA LIPs). The suspensions were then subjected to probe sonication using a tip sonicator (Branson SFX250, USA) for 6 cycles of 10 seconds each at 10% amplitude. All liposome preparations were dialyzed for 24 h using 6-8 kDa dialysis membranes to remove unencapsulated ICG. Each liposome formulation was divided into two aliquots: one fraction was centrifuged (Eppendorf, 5424R, Germany) at room temperature for 15 min at 20,000 g (Cent +), while the other one was kept without centrifugation (Cent -).

**Table 1.** Lipid ratios and amounts used for the preparation of ICG-PEG and ICG-HA LIPs.

| Liposomes | Lipid composition<br>*Lipid ratios* | m<sub>DOTAP</sub> | m<sub>DSPC</sub> | m<sub>DSPE-PEG2kDa</sub> | Rehydration medium |
| --- | --- | --- | --- | --- | --- |
| ICG-PEG LIPs | DOTAP/DSPC/DSPE-PEG2kDa<br>50/47.5/2.5 | 1.67 mg | 1.79 mg | 0.335 mg | 1 ml water |
| ICG-HA LIPs | DOTAP/DSPC<br>50/50 | 1.67 mg | 1.89 mg | --- | 1 ml 2.5mg/ml DOPE-HA in water |

**Scheme 1.**
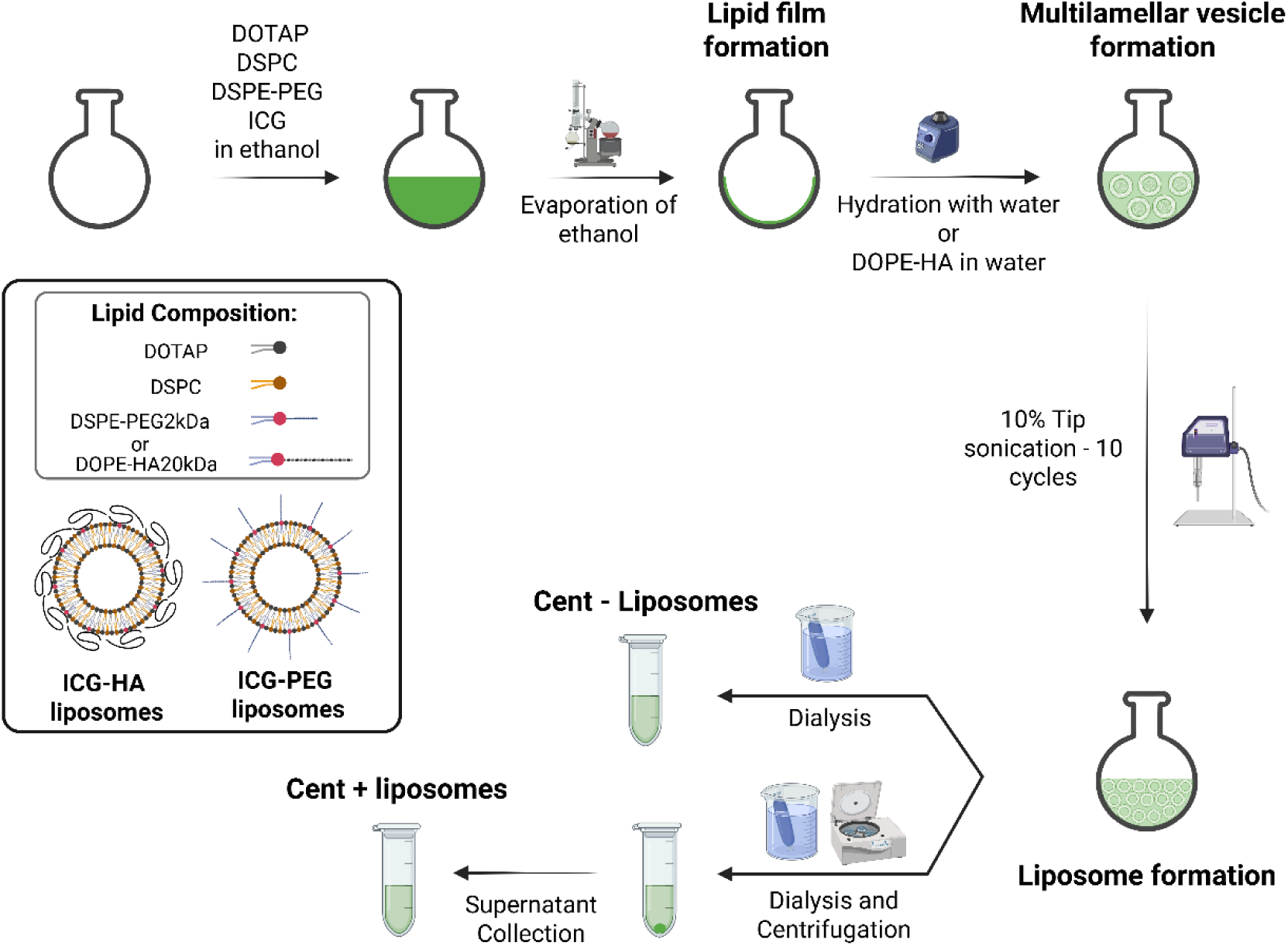
Preparation of PEG-coated and HA-coated ICG liposomes via the thin lipid film rehydration method. At the end of the preparation process, two liposome formulations were obtained; Cent + liposomes, which were dialyzed and then centrifuged, and Cent - liposomes that were only dialyzed. Image created with BioRender.com.

The liposome size was characterized by dynamic light scattering (DLS) (Malvern Panalytics, USA) before and after centrifugation. Zeta potential was measured by DLS but only for the centrifuged LIPs. Encapsulation efficiency (EE%) was measured for both Cent + and Cent - LIPs by quantifying ICG concentration using UV-Vis spectrophotometry (NanoDrop2000, ThermoFisher Scientific, USA).

### Preparation and characterization of HA-coated ICG nanoaggregates

Scheme 2 summarizes the preparation of ICG nanoaggregates (ICG AGG NPs). In brief, a 5 ml aqueous solution of synthesized DOPE-HA (0.25 mg/ml, equivalent to 0.0125 mM) was prepared. Then, 500 µL of a 1 mg/ml ICG powder dispersion in chloroform was added dropwise to the DOPE-HA solution under continuous stirring, while applying probe sonication at 10% amplitude for 6–10 cycles (10 seconds per cycle, with 5-second intervals). ICG did not dissolve in chloroform but was only dispersed. To micronize the ICG powder, the dispersion was sonicated in a water bath (Branson 2800Mh, USA) for 15 minutes prior to use. After addition, the mixture was stirred at 250–300 rpm (Heidolph MR 3001K, Germany) for 12–24 h to ensure complete chloroform evaporation. The resulting ICG AGG NPs were purified by centrifugation at 20,000 g for 15 minutes and resuspended in water at the desired concentration.

**Figure 2.**
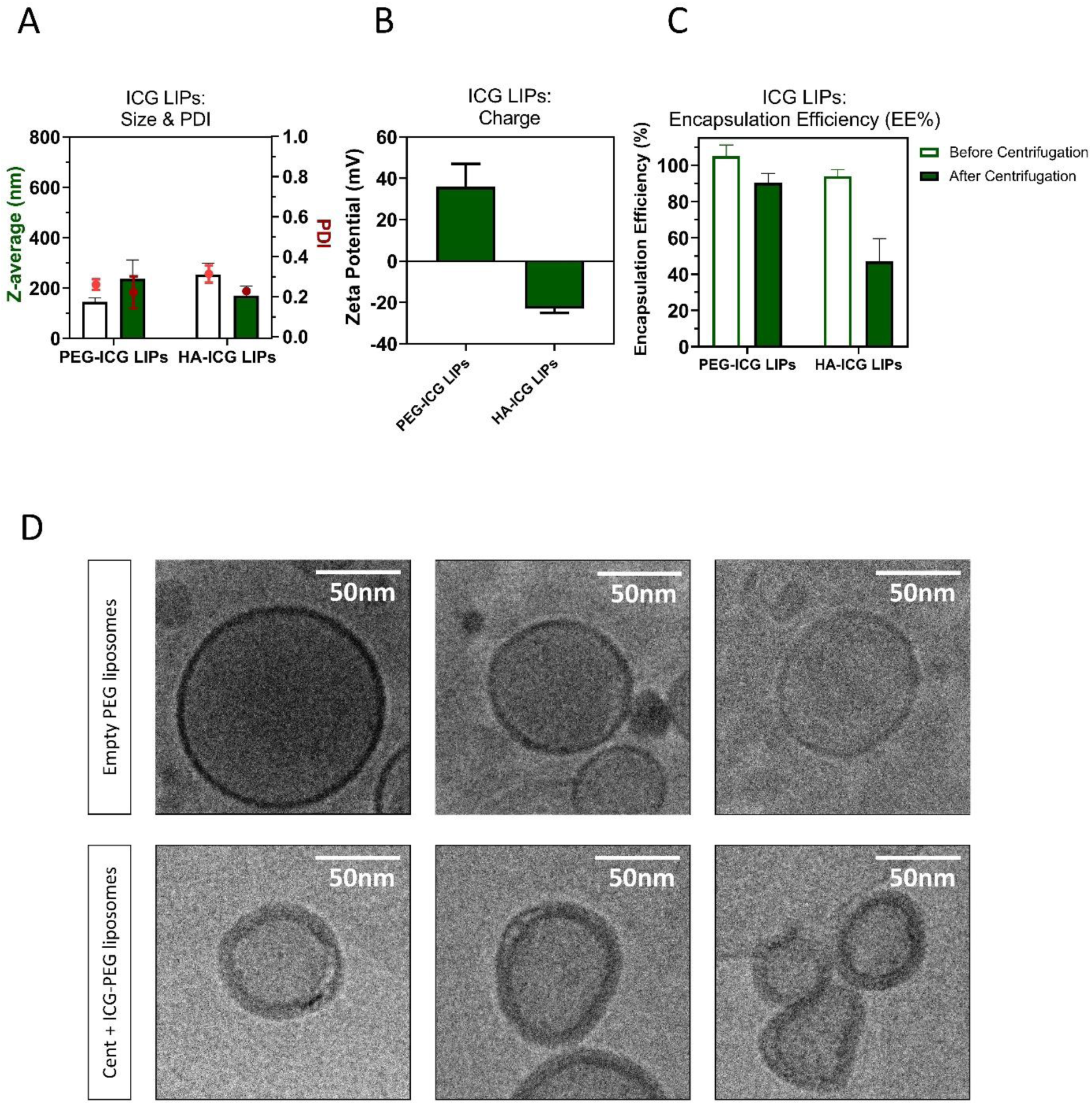
Characterization of the ICG liposomes. **A)** Size and polydispersity (PDI) and **B)** zeta potential of ICG-loaded liposomes coated with PEG or HA characterized by DLS (n= 3). **C)** Encapsulation efficiency of ICG-PEG and HA-ICG LIPs before (Cent -) and after (Cent +) centrifugation (20000 g) measured by UV/Vis spectrophotometry (n= 3). **D)** Representative cropped cryo-EM micrographs (n= 3) of empty PEG-coated liposomes (control) and Cent + ICG-PEG liposomes. Scale bar 50 nm.

The size and charge of the prepared ICG AGG NPs were characterized by DLS (Malvern Panalytics, USA). UV-Vis absorption measurements (Shimadzu, Japan) of ICG AGG NPs were performed in water and compared with free ICG at an equivalent concentration of 32 µg/ml. ICG AGG NP fluorescence (Perkin Elmer, USA) was measured compared to free ICG at a 800 nm emission wavelength using a 780 nm excitation wavelength in a 96 well plate at 6.25, 12.5, 25 and 50 µg/ml ICG concentrations.

**Scheme 2.**
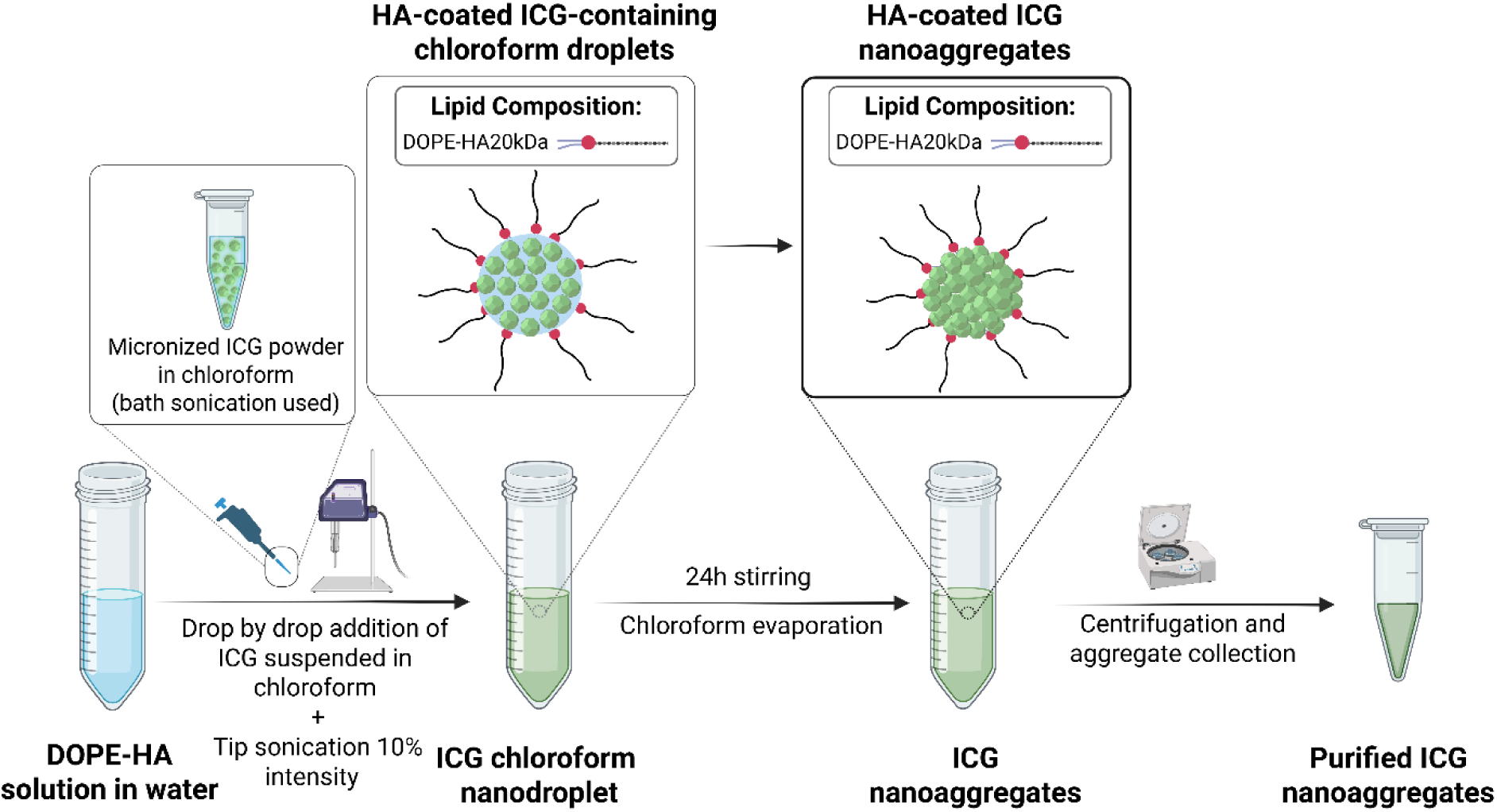
The method employed for the preparation of ICG nanoaggregates coated with HA. This method was inspired by the emulsification-solvent evaporation technique and modified by using an undissolved dispersion of ICG in chloroform instead of solubilized ICG. Image created with BioRender.com.

### VNB generation characterization of liposomes and nanoaggregates

VNB generation was observed in microscopy dishes (MatTek Corporation, USA). Dark-field microscopy was used for imaging, as VNBs scatter light and appear as bright spots [34]. Samples were illuminated with nanosecond laser pulses at a wavelength of 532 nm with a pulse duration <7 ns (Ekspla N230-100-SCU-2H OPO) and repetition rate of 100 Hz. Either 561 nm or 800 nm wavelengths were used for these experiments. For the 561 nm laser wavelength, a beam expander (#GBE05-A, Thorlabs) combined with an iris diaphragm (#D37SZ, Thorlabs) was used to adjust the diameter of the laser beam spot to 160 μm. For experiments carried out at 800 nm, the diameter of the laser spot was adjusted to 114 µm. The energy delivered by the laser at both wavelengths was monitored by an energy meter (J-25MB-HE&LE, Energy Max-USB/RS sensors, Coherent) synchronized with the pulsed laser.

40 µl samples of Cent + and Cent - ICG-PEG LIPs containing 0.56 mg/ml and 0.66 mg/ml ICG were transferred into the microscopy dishes and irradiated with 2.5 and 2.6 J/cm^2^ single laser pulses, respectively. Dark-field images were acquired before and during illumination with a single <7 ns laser pulse. For this experiment, only the 561 nm wavelength was used for PS excitation. The resulting images were then used to visualize VNBs.

VNB generation by ICG AGG NPs was analyzed in the same manner as for LIPs. For these experiments, ICG AGG NPs at a concentration of 0.033 mg/ml ICG were used. VNB formation was visualized at both 561 nm and 800 nm with laser fluences of 1.5 and 3.2 J/cm^2^, respectively.

### VNB threshold calculations

For the determination of the VNB generation threshold for each formulation, the number of visible VNBs within the laser spot area was quantified at laser fluences ranging from 0.2-5.2 J/cm^2^. The VNB threshold, defined as the laser fluence at which at 90% of the irradiated NPs generate VNBs [34], was calculated by fitting the VNB count as a function of laser fluence using a Boltzmann sigmoid model.

### Electron cryo-microscopy (cryo-EM)

These experiments were performed on ICG AGG NPs, Cent + ICG PEG LIPs, and PEG LIPs without ICG (control -). ICG AGG NPs had an ICG concentration of 0.33 mg/ml. Cent + ICG-PEG LIPs and PEG LIPs without ICG were concentrated 5 times compared to the original preparation, using Amicon Ultra 100kDa centrifugal filters, to obtain a lipid concentration around 20 mg/ml for better imaging.

For the empty PEG LIPs and Cent + ICG-PEG LIPs samples, 4-µl samples were applied to glow-discharged R2/1 300 mesh holey carbon copper grids (Quantifoil Micro Tools GmbH). For the ICG AGG NPs, 4 µl of a sample was applied to glow-discharged Lacey Formvar/Carbon 400 mesh copper grids (Ted Pella, Inc.). Grids were blotted and plunge frozen in liquid ethane using a Lecia EM GP2 Plunge Freezer (Leica Microsystems) operated at 95% humidity at 20°C. Due to the dye present in the ICG-PEG LIPs and ICG AGG NPs, sensor blotting was not possible, and therefore a manually chosen move of 44 mm was used with 4 sec blot time for both samples. For the empty PEG LIPs, sensor blotting was performed using 2 mm additional move and 4 sec blot time. Micrographs were recorded at the VIB Bioimaging Core Ghent (Zwijnaarde, Belgium), on a JEOL 1400+ TEM operated at 120 keV, equipped with a JEOL Ruby CCD camera. Images were taken at a magnification of 60.000X, resulting in a pixel size of 2.639 Å/pixel at the specimen level.

### Preparation of the artificial collagen fibers and the collagen cloud model

In this study, collagen type I was destabilized to either form thread-shaped fibers resembling vitreous opacities or to form cloud-like structures used for the quantification of photodisruption. Thread-shaped collagen fibers were prepared as previously described, yielding a final collagen concentration of 0.2 mg/ml [15]. To obtain the collagen cloud model, a 0.2 mg/ml dispersion of thread-shaped fibers was sonicated at 10% intensity for 5 min prior to use in the quantification experiments.

### Qualitative inspection of collagen fiber photodisruption

To investigate VNB-mediated photodisruption, the ability of ICG-PEG LIPs and ICG AGG NPs to photodisrupt collagen fibers was analyzed by dark-field microscopy. All samples were placed into microscopy slides (Marienfeld, Germany). Fibers incubated with ICG-PEG LIPs were irradiated at 561 nm. Cent + and Cent - ICG-PEG LIPs contained ICG at final concentrations of 0.56 mg/ml and 0.66 mg/ml, respectively. In comparison, fibers incubated with ICG AGG NPs were irradiated at both 561 nm and 800 nm, with a final ICG concentration of 0.43 mg/ml. All groups were incubated with collagen fibers at 0.02 mg/ml for 24 h at room temperature prior to laser irradiation (Ekspla N230-100-SCU-2H OPO). To improve imaging, the original collagen fiber dispersion (0.2 mg/ml) was diluted 10-fold to increase inter-fiber spacing [15].

### Quantification of PS photodisruption using the collagen cloud model

Photodisruption was quantified using a collagen cloud model consisting of randomly dispersed collagen structures at a final concentration of 0.2 mg/ml. Quantification was performed in five groups: ICG AGG NPs containing 0.33 mg/ml ICG, both cent + ICG-PEG and ICG-HA LIPs containing 0.46 mg/ml and 0.56 mg/ml ICG, respectively, a free ICG solution at 0.56 mg/ml, and a laser-only control group without PS. The collagen cloud was incubated with each formulation for 24 h at room temperature. Subsequently, 40 µL of each sample was transferred into 50 mm glass-bottom dishes (MatTek Corporation, USA) and irradiated with the pulsed-laser (Ekspla N230-100-SCU-2H OPO) at 561 nm (spot size: 160 µm, pulse duration ˂7ns).

Data acquisition was performed by capturing dark-field microscopy images of each sample before laser exposure (*II_before pulse_*) and after each laser pulse (*II_pulse n_*). To ensure that scattering intensity reflected only the collagen cloud and not the PS formulations, control images were acquired from each formulation in the absence of collagen (Figure S6). These control images were used to determine the background scattering intensity (*II₀*) specific to the PS alone. The *II₀*values were subtracted from the measured intensities of collagen-containing samples (before and after laser exposure) to isolate the collagen-specific scattering signal. The percentage of scattering intensity within the irradiated area was then quantified using Fiji ImageJ software by measuring the integrated pixel density inside the laser beam spot, according to Equation 1:

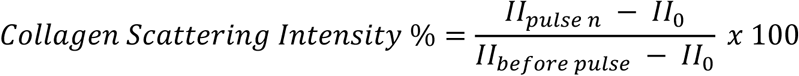

***Equation 1. Calculation of scattering intensity % to quantify photodisruption in the collagen cloud.*** *Parameters: II_pulse n_ (integrated intensity in the beam area for PS-incubated collagen after n laser pulse(s)), II_before pulse_ (integrated intensity in the beam area for PS-incubated collagen before irradiation) and II_0_ (integrated intensity in the beam area of the tested PS alone (without collagen))*.

All experimental settings—including the position of the dark-field condenser relative to the sample, illumination intensity, and optical filters used during image acquisition—were kept constant across all groups to ensure consistency and accuracy in quantification.

### Toxicity studies in retinal cells in vitro

To assess the potential cytotoxicity of ICG AGG NPs compared with free ICG, a CellTiter-Glo® viability assay (Promega, USA) was performed on ARPE-19 and MIO-M1 cell lines, representing retinal pigment epithelial (RPE) and Müller glial cells, respectively. ARPE-19 cells (ATCC, Maryland, USA) were cultured in DMEM/F-12 medium supplemented with 10% fetal bovine serum (Hyclone, Cramilton, UK), 1% L-glutamine (Gibco, Paisly, UK) and 2% penicillin-streptomycin (Gibco, Paisly, UK). MIO-M1 cells were obtained from the UCL Institute of Ophthalmology (London, UK) and cultured in DMEM GlutaMAX with low glucose (Gibco, Paisly, UK) supplemented with 10% fetal bovine serum (Hyclone, Cramilton, UK), 1% L-glutamine (Gibco, Paisly, UK) and 2% penicillin-streptomycin (Gibco, Paisly, UK). For each cell type, experiments were performed in triplicate. Both cell types were seeded at a density of 10⁴ cells per well in a 96-well plate (Greiner, Austria). After 24 h, cells were incubated with either free ICG or ICG AGG NPs at five concentrations ( 0-50 µg/ml ICG), prepared in FBS-free medium, for 4 h at 37°C in 5% CO_2_. Each concentration was tested in quadruplicate.

After incubation, treatment solutions were removed and cells were washed with 100 µl PBS (-/-). Following PBS removal, 100 µl of CellTiter-Glo® 2.0 reagent diluted 1:1 in PBS was added to each well. The plate was then placed on an orbital shaker (Interteck, UK) at 80 rpm for 5–15 min. Fluorescence was then measured using a GloMax® plate reader (Promega, USA) with excitation at 485–500 nm and emission at 520–530 nm. Cell metabolic activity was expressed as a percentage of the average signal from untreated controls (PBS only).

## Results

### Preparation and physicochemical characterization of ICG-loaded liposomes

The ICG-loaded liposomes were formulated by the thin-film rehydration method as illustrated in Scheme 1 and similar to previous literature reports [33]. For all prepared liposomes, the main lipid components were the cationic lipid DOTAP and DSPC. These liposomes were coated either with hyaluronic acid (HA) or polyethylene glycol (PEG). The rationale behind the use of DOTAP was to harness its positive charge to allow for electrostatic interactions with the negatively charged sulfonic groups of the ICG molecules. By doing so, we expect (i) to improve ICG encapsulation efficiency (EE%) thanks to the affinity of ICG to DOTAP [35], and ii) to facilitate the coating of the liposomes with HA which is negatively charged. After liposome preparation, all formulations were dialyzed for 24h. The dialyzed formulations were then divided into two groups. One group was subjected to highspeed centrifugation (Cent +) to remove the dye aggregates that cannot be dialyzed, while the other was directly collected after dialysis without centrifugation (Cent -).

Both HA-coated and PEGylated liposomes, namely ICG-HA LIPs and ICG-PEG LIPs, exhibited similar sizes before and after centrifugation (Figure 2A). PDI values for both particle types, before and after centrifugation, remained comparable, ranging from 0.2 to 0.3 (Figure 2A). The results obtained for Cent + and Cent - ICG-HA and PEG LIPs indicate that in Cent - formulations, either liposomes had a similar size to the aggregates or that aggregation could not be detected at all by DLS. Measuring the NP surface charge showed that ICG-HA LIPs were negatively charged, confirming the presence of HA at the surface of the LIPs, while ICG-PEG LIPs remained positively charged due to the presence of (neutral) PEG on their surface (Figure 2B). Finally, the EE% measurements indicated that there was no free ICG in both Cent - ICG-HA and ICG-PEG LIPs, as the EE% of these NPs after dialysis and before centrifugation was around 100% for both groups. However, after centrifugation (Cent +), the EE% decreased by 15% for ICG-PEG LIPs and by 50-60% for ICG-HA LIPs (Figure 2C).

Given the observed accumulation of sediments and the drop of EE% after centrifugation, we hypothesize the presence of two distinct NP populations in the formulations prior to centrifugation. The first population likely consists of liposomes that remained in the supernatant post-centrifugation and encapsulated approximately 85% and 40-50% of ICG inside ICG-PEG LIPs and ICG-HA LIPs, respectively. The second population likely consists of high-density NPs that sedimented upon centrifugation, likely due to significant ICG aggregation—either in pure form or occurring within the liposomes.

Given the higher EE% of ICG-PEG LIPs compared to ICG-HA LIPs after centrifugation, which indicates a better encapsulation of ICG, we conducted cryo-EM imaging on Cent + ICG-PEG LIPs and compared them with empty PEG liposomes (no ICG) as the control. As shown in Figure 2D, top panel, empty PEG LIPs display a hollow core corresponding to the aqueous core surrounded by a thin bilayer. In contrast, ICG-PEG LIPs exhibit a much thicker bilayer (Figure 2D, bottom panel), indicative of ICG incorporation in the bilayer while the core remains hollow.

### VNB generation characterization of ICG-loaded liposomes

To determine the capacity of ICG-loaded liposomes to induce VNBs, we evaluated the response of Cent + and Cent - ICG-PEG LIPs to laser irradiation (<7 ns; 561 nm) by visualizing VNB generation using dark-field microscopy. ICG-PEG LIPs were chosen over ICG-HA LIPs for these experiments due to their significantly higher EE% after centrifugation, indicating a higher ICG encapsulation.

Figure 3A shows a dark-field microscopy image of Cent - ICG-PEG LIPs (0.66 mg/ml ICG). As shown in this figure, upon a single nanosecond laser pulse (2.6 J/cm^2^), the formulation exhibited multiple bright spots during irradiation, corresponding to the formation of VNBs. In contrast, no VNBs were detected using Cent + ICG-PEG LIPs with similar laser settings (2.51 J/cm^2^) (Figure 3B). The number of VNBs generated by Cent - ICG-PEG LIPs was counted for increasing laser pulse fluence. From this, we found that the VNB threshold —defined as the laser pulse fluence at which 90% of the irradiated NPs generate VNBs [34]— was 1.1 J/cm^2^ (Figure 3C).

**Figure 3.**
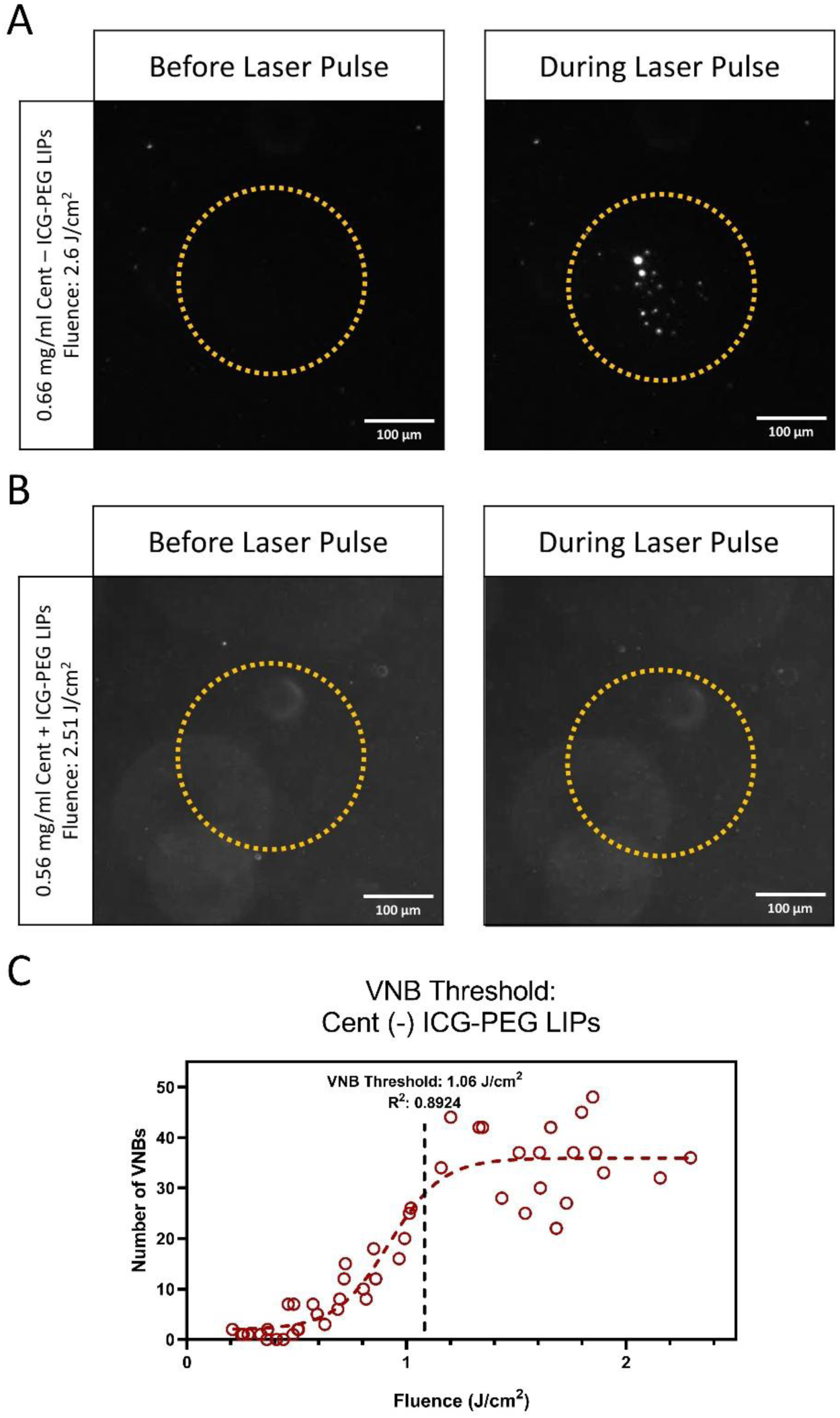
Reactivity of ICG-PEG LIPs to pulsed-laser irradiation (<7ns; 561 nm). Representative dark-field images of **A)** Cent - ICG-PEG LIPs and **B)** Cent + ICG-PEG LIPs before and during illumination with a single nanosecond laser pulse (561 nm). Scale bars for all dark-field images: 100 µm. **C)** The VNB threshold of Cent - ICG-PEG LIPs was calculated by fitting the number of generated VNBs as a function of the fluence to the Boltzmann sigmoid curve.

This observation supports the hypothesis of the existence of a second population of particles in the Cent - formulation which likely corresponds to aggregated ICG and is more likely to generate VNBs.

### In vitro evaluation of floater photodisruption by liposomes

To further investigate the efficacy of these liposomal formulations for the photodisruption of vitreous opacities, artificial collagen fibers were incubated with either Cent + or Cent - ICG-PEG LIPs, irradiated by laser pulses and observed by dark field microscopy.

As shown in Figure 4A (Movie S1), Cent - lCG-PEG LIPs (0.66 mg/ml ICG) exhibited substantial collagen fiber photodisruption already after four laser pulses of ∼1 J/cm². However, Cent + ICG-PEG LIPs (Figure 4B, Movie S2) did not induce any fiber photodisruption, even after 20 pulses of 2.3 J/cm^2^. The outcome with the Cent + liposomes remained comparable to the fibers exposed to the laser in the absence of any PS (Figure 4C, see Movie S3). Indeed, pulsed-laser application on fibers incubated with this formulation induced only limited photomechanical effects, resulting in slight fiber displacement but no noticeable photodisruption.

**Figure 4.**
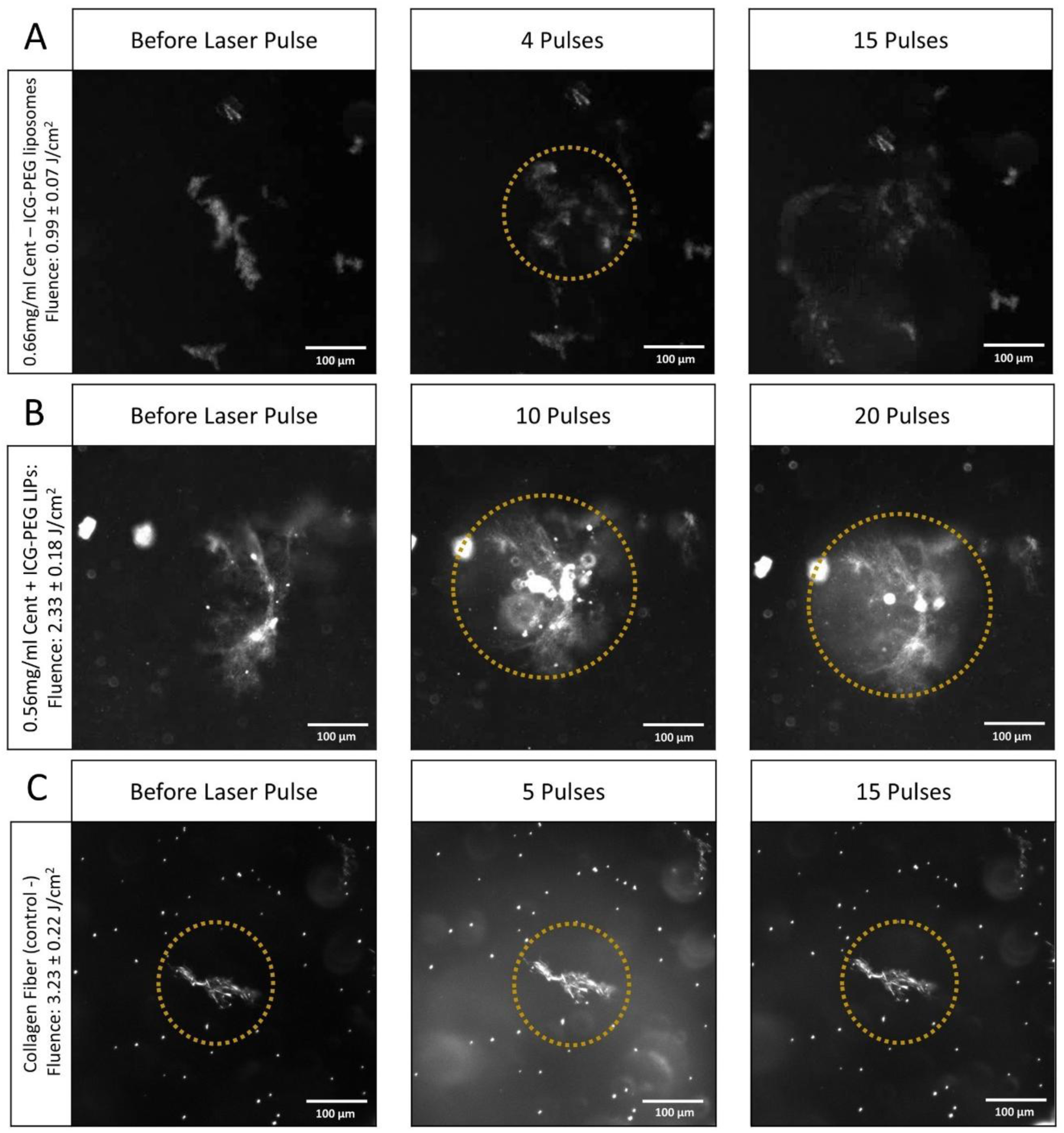
**In vitro collagen fiber photodisruption assessment of ICG-PEG LIPs before and after 561 nm nanosecond pulsed-laser irradiation using dark-field microscopy**. Dark-field microscopy images were obtained from **A)** Cent - and **B)** Cent + ICG-PEG LIPs versus **C)** only collagen fibers (no PS) before and after a certain number of laser pulses. Scale bar 100 µm.

Overall, these results highlighted that ICG aggregates separated from or within the liposomes were likely responsible for photodisruption. Indeed, only the Cent - formulation, in which two populations of ICG particles were suspected, exhibited photodisruption. On the upside, this experiment confirmed the critical role of VNBs for inducing photomechanical forces for effective collagen fiber photodisruption, as shown previously by our group with gold nanoparticles [15].

### Preparation and photophysical characterization of ICG nanoaggregates

The pronounced photodisruption observed with Cent - ICG-PEG LIPs was likely due to the presence of ICG aggregates, either associated with the liposomes or existing freely in the dispersion. Therefore, we next investigated ICG nanoaggregates as a potential approach for the photodisruption of vitreous opacities. To accomplish this, we designed ICG aggregated NPs (ICG AGG NPs), composed of aggregated ICG and using HA-conjugated DOPE as a stabilizing agent (Scheme 2). To prepare ICG AGG NPs, we employed a similar approach to the emulsification-solvent evaporation technique, as reported in the literature [36, 37].

ICG AGG NPs were characterized using DLS, finding a hydrodynamic diameter of 463 ± 76 nm (Figure 5A) with a negative zeta potential of -25.0 ± 1.4 mV (Figure 5B), similar to ICG-HA LIPs. Interestingly, these NPs stored in water at 4°C remained stable in size and PDI for two months (Figure S2A,B). Moreover, the successful formulation of ICG AGG NPs was supported by multiple lines of evidence: (i) comparison of the light absorption spectra of free ICG and ICG AGG NPs (at equivalent ICG concentrations) revealed a reduced peak intensity for ICG AGG NPs, accompanied by extended absorption tails on both sides of the spectrum (Figure 5C). This spectral shift indicates disordered aggregation [38], in contrast to the ordered H- and J-aggregates of ICG, which display sharp absorption peaks around 700 nm and 890 nm, respectively [39, 40]; (ii) ICG fluorescence was markedly quenched upon formulation into ICG AGG NPs (Figure 5D), consistent with the aggregation-caused quenching (ACQ) effect [17]; and (iii) cryo-EM imaging demonstrated that ICG AGG NPs are spherical particles with a highly electron-dense core, consistent with ICG accumulation (Figure 5E).

**Figure 5.**
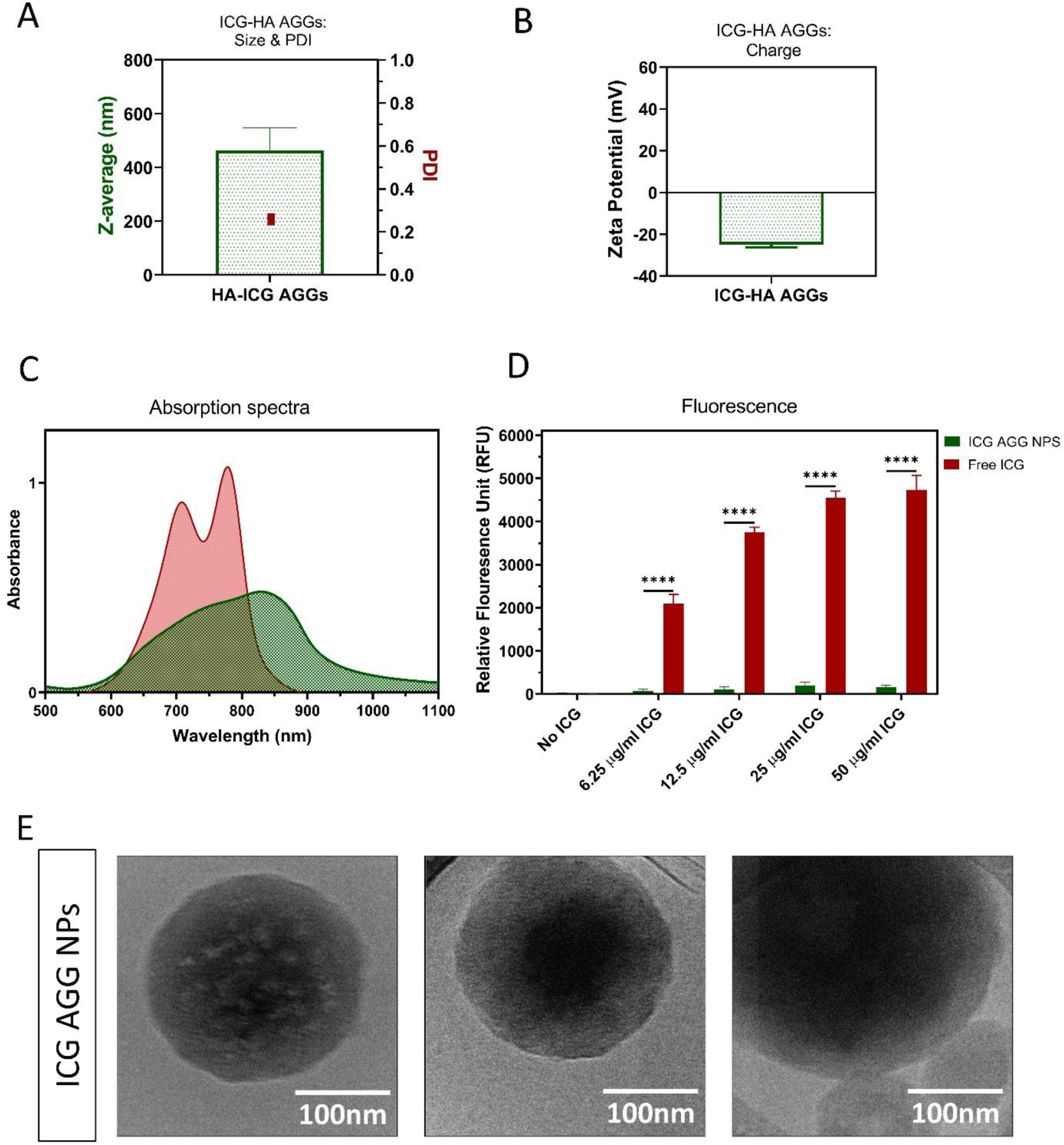
Characterization of ICG AGG NPs. **A)** Size and polydispersity (PDI) and **B)** zeta potential of ICG AGG NPs coated with HA (n= 3). **C)** Absorption spectra and **D)** fluorescence of free ICG (red) and ICG AGG NPs (green) (λ Ex/λ Em 780/800 nm). Unpaired t-test **** P ˂ 0.0001. **E)** Representative cropped cryo-EM images (n= 3) of ICG AGG NPs. Scale bar 100 nm.

Finally, we showed that the use of DOPE-HA as a stabilizer is crucial for the successful preparation of ICG AGG NPs. We were able to confirm that DOPE-HA promotes the formation of monodisperse ICG AGG NPs with high particle counts (Figure S3). As shown in Figure S3A, using DOPE-HA during the preparation resulted in NPs with a PDI of 0.2-0.3 and a high derived count rate of approximately 4000. The derived count rate reflects the total intensity of scattered light and serves as an indirect measure for the number of NPs present within the sample. Interestingly, in the absence of DOPE-HA, a broader PDI and a substantially lower derived count rate (∼150 ) was observed, suggesting that most of the ICG was dissolved in water, and that the few remaining aggregates were highly heterogeneous. In addition, visual inspection of both stabilized and non-stabilized formulations supported this conclusion: the DOPE-HA stabilized formulation exhibited a darker green color, indicative of a higher concentration of ICG nanoaggregates (Figure S3B).

### Characterization of VNB generation and in vitro floater photodisruption by ICG nanoaggregates

To examine the capacity of ICG AGG NPs to generate VNBs and the effect of the excitation wavelength, they were irradiated at 561 nm, similar to liposomes, and at 800 nm, which is around the excitation wavelength maximum of these NPs. Visual inspection of VNB formation indicated that ICG AGG NPs successfully generated VNBs upon laser irradiation at both laser wavelengths, as evidenced by the appearance of bright spots visible in dark-field microscopy images (Figure 6A, left panel). Furthermore, the VNB thresholds for both wavelengths were found to be around 1.78 J/cm^2^ at 561 nm and 1 J/cm² at 800 nm (Figure 6A, right panel). Although the threshold was lower at 800 nm, both fluences can be considered comparable when viewed in the context of typical values used in clinical practice (Figure 6A, right panel). Taken together with the light absorption properties of ICG AGG NPs at 561 nm and 800 nm wavelengths (Figure 5C), these findings indicate that absorption does not play a critical role in the generation of VNBs in our experimental conditions.

**Figure 6.**
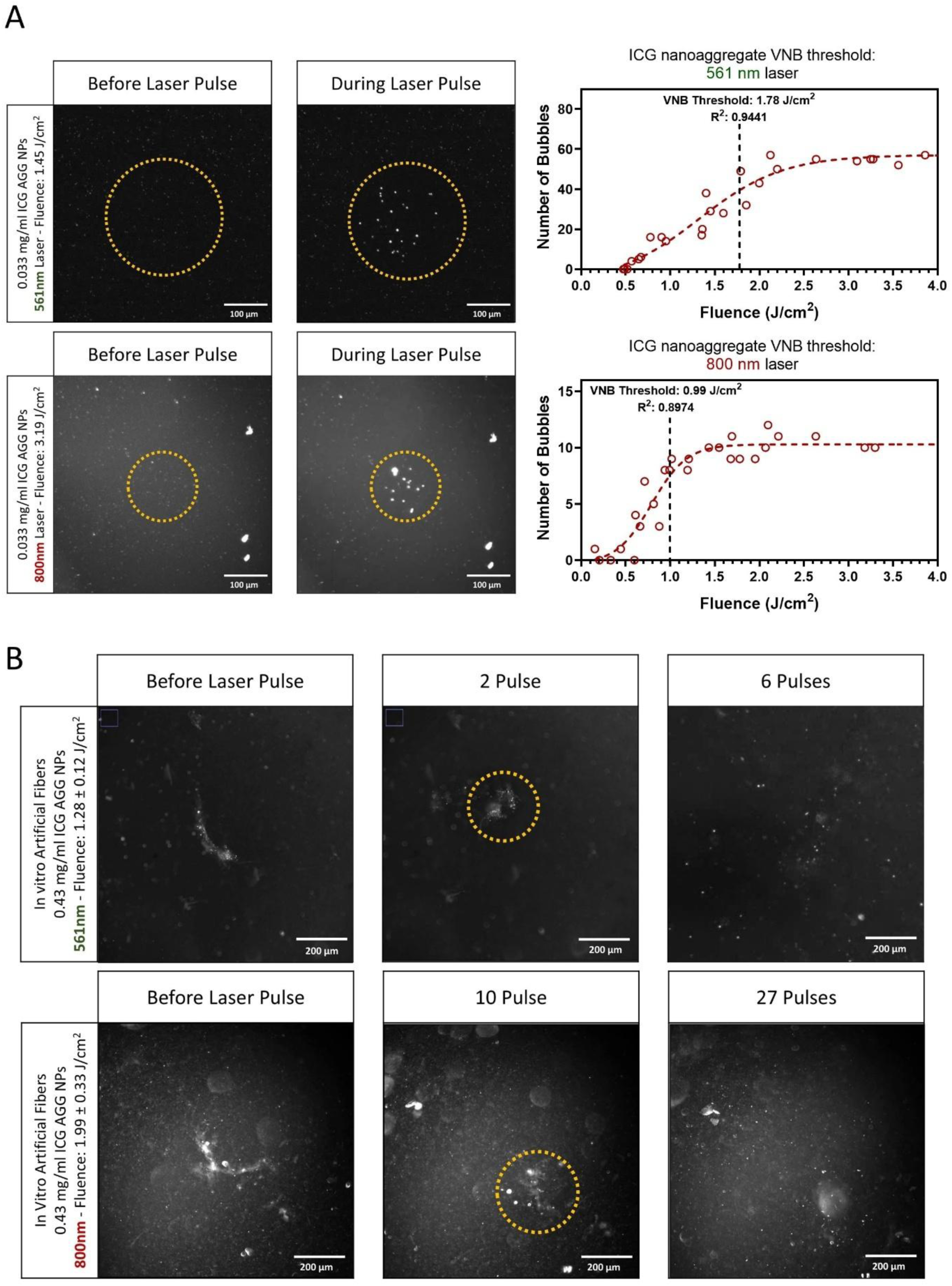
VNB formation and in vitro fiber photodisruption of ICG AGG NPs upon irradiation with 561 nm and 800 nm nanosecond laser pulses. **A)** The VNB formation of ICG nanoaggregates at 561 nm and 800 nm was assessed by dark-field microscopy (scale bar 100 µm). VNB thresholds at both wavelengths were determined by fitting the number of generated VNBs as a function of fluence to a Boltzmann sigmoid curve. **B)** The ability of ICG nanoaggregates to destroy collagen fibers in vitro at 561 nm (1.28 j/cm^2^) and 800 nm (1.99 j/cm^2^), was assessed by dark field microscopy. Scale bar 200 µm.

After determining the capacity for VNB formation, we assessed the photodisruption efficacy of ICG AGG NPs on collagen fibers at both 561 nm and 800 nm and at fluences around the calculated VNB threshold to ensure sufficient VNB generation. The top and bottom panels of Figure 6B show representative images of fibers that were incubated with ICG AGG NPs (0.43 mg/ml ICG) and treated using 561 nm (Figure 6B, top panel; see also Movie S4) and 800 nm (Figure 6B, bottom panel; see also Movie S5) nanosecond laser pulses with fluences of 1.3 J/cm^2^ at 561 nm and 2.0 J/cm^2^ at 800 nm. Using laser pulses at both wavelengths, fibers underwent complete disruption. In contrast, in control experiments in which fibers were exposed to laser light in the absence of ICG AGG NPs, no noticeable disruption occurred at 561 nm (Figure 4C; see Movie S3) and 800 nm (Figure S4; see Movie S6) laser wavelengths. These findings again confirm a strong correlation between VNB formation and photomechanical fiber disruption.

Given the difference between the results obtained with ICG AGG NPs and ICG LIPs, it became evident that the ability of ICG to generate VNBs is linked to supramolecular aggregation. A likely explanation of the absence of VNB is the absence of ICG clustering within the liposomal bilayer and within the core as seen with the lack of contrast in the core, as opposed to ICG AGG NPs exhibiting-denser cores (Figure 5E). This densely packed core of ICG AGG NPs may enhance VNB generation by increasing the NP’s thermal capacity, thereby facilitating heat accumulation, leading to more intense vapor formation and subsequently, stronger mechanical forces.

### In vitro model for quantification of floater photodisruption

Visual examinations of individual fibers confirm the occurrence of photodisruption but do not provide quantitative data. In principle, one could quantify the number of pulses required to disrupt a defined number of fibers as a metric for comparing the photodisruption efficacy of different formulations. However, the inherent heterogeneity in fiber size, shape and collagen density makes it challenging to draw standardized conclusions from such measurements.

Therefore, rather than looking at individual fibers, we propose to make use of a relatively-homogenized suspension with a high concentration of fibers. When imaged by dark field microscopy, light is scattered by the fibers, resulting in a certain image brightness. When fibers are destroyed, light scattering becomes less efficient and results in a decrease in the observed intensity. This relative decrease in scattered light intensity can be considered a measure for fiber photodisruption. An example is shown in Figure 7A for ICG-PEG LIPs and ICG AGG NPs. Here, a collagen fiber suspension (0.2 mg/ml) was sonicated until a visually uniform cloud-like dispersion was formed. This fiber suspension was then incubated with the formulation of interest for 12-24 h and imaged by dark-field microscopy before and after applying a number of laser pulses (1, 4 and 8 pulses). As the images show, the more pulses are applied, the more the scattered light intensity decreases within the irradiated area for ICG AGG NPs whereas for Cent + ICG-PEG LIPs, scattered light intensity remains unchanged throughout laser irradiation.

**Figure 7.**
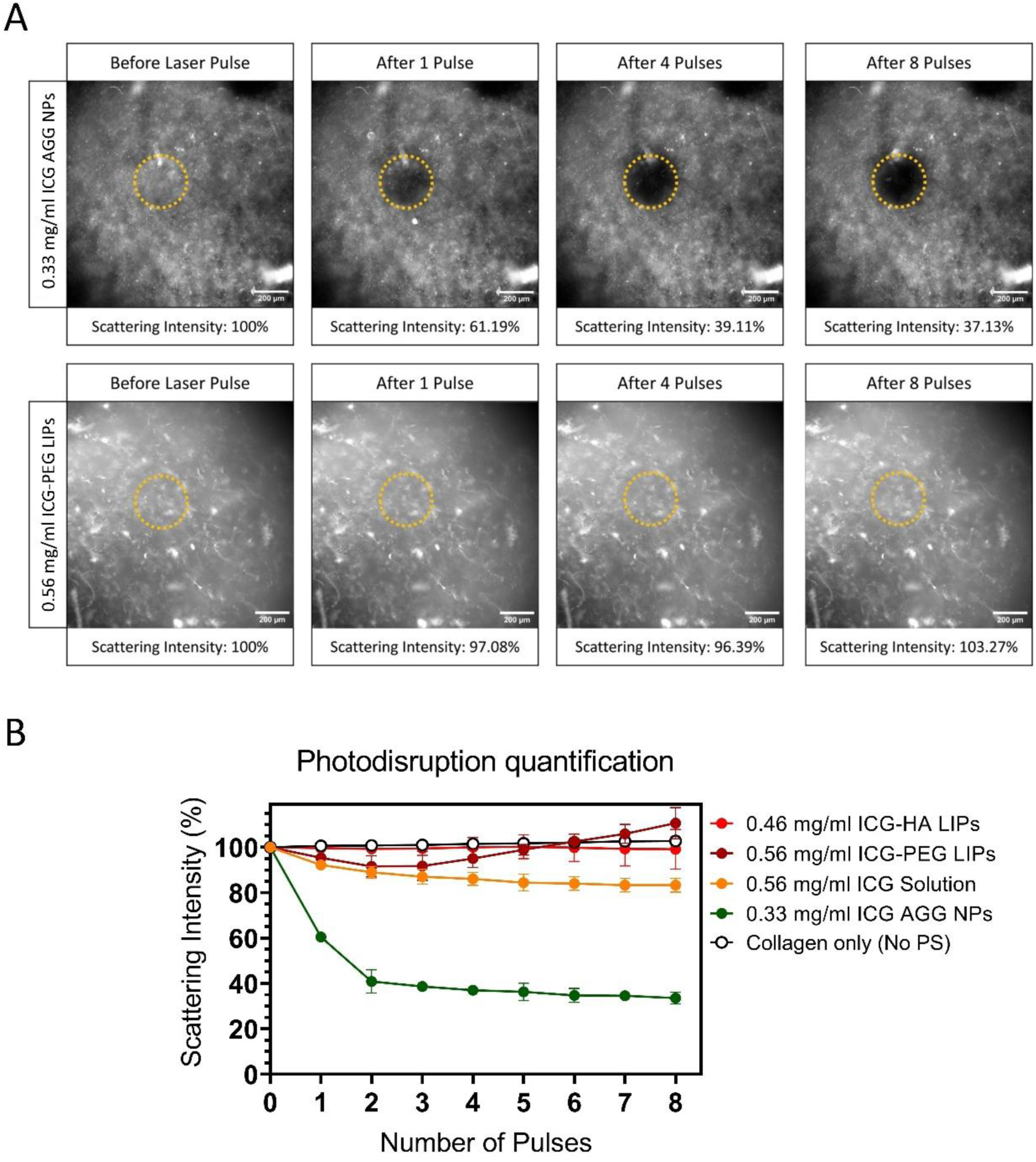
Quantification of collagen fiber photodisruption upon nanosecond laser irradiation at 561 nm. **A)** Representative dark-field microscopy images used for photodisruption quantification of ICG AGG NPs vs ICG-PEG LIPs. Scale bar 200 µm. Representative images of other tested conditions can also be found in the supplementary information in Figure S5. **B)** Relative scattering intensity data obtained from these groups (n= 4 per group).

In total, five groups were tested in these experiments: ICG AGG NPs containing 0.33 mg/ml ICG, both cent + ICG-PEG and ICG-HA LIPs containing 0.46 mg/ml and 0.56 mg/ml ICG, respectively, free ICG at 0.56 mg/ml, and a control group exposed to laser only (no PS). ICG-HA LIPs were included for this set of experiments for a direct comparison with ICG AGG NPs, as both are coated with HA. Indeed, since HA is expected to maintain affinity towards collagen fibers [15], one might argue that ICG-HA LIPs are more effective compared to ICG-PEG LIPs due to a better accumulation and aggregation on collagen fibers. All PS groups were incubated with fiber samples for 24 h. Following incubation, laser treatment was applied, and scattering intensity was monitored for up to 8 pulses at 561 nm and a fluence of ∼3 J/cm^2^. Representative dark-field images for free ICG, ICG-HA LIPs, and collagen-only (no PS) groups are shown in Figure S5.

The relative scattered light intensity for each group is presented in Figure 7B. Irradiation of the fiber suspension in the absence of NPs was ineffective. In contrast, ICG AGG NPs induced up to a 70% reduction in scattered light, demonstrating efficient photodisruption already after 2 to 3 pulses while free ICG only resulted in around 20% reduction of the scattered light, even after 8 pulses. We attribute this observation to the fact that free ICG must first aggregate at the surface of the collagen in order to generate VNBs. However, the ICG aggregation / binding process is rather limited and poorly controlled. In contrast, ICG AGG NPs are pre-formed, homogeneously aggregated structures that are expected to be capable of more efficient and abundant VNB generation. Both ICG-HA LIPs and ICG-PEG LIPs showed no photodisruption effect, confirming that ICG encapsulation in liposomes does not enable VNB-mediated photodisruption.

### In vitro safety evaluation of ICG nanoaggregates

Despite the anticipated overall improvement in the retinal safety of ICG NPs compared to free ICG, there remains a risk that a portion of these NPs could penetrate specific parts of the retina, causing toxicity. Moreover, ICG AGG NPs are coated with HA, which could further contribute to toxicity by enhancing NP cellular uptake through interactions between HA and CD44 receptors expressed by retinal cells such as Müller glia and retinal pigment epithelium (RPE) cells [41–43]. Therefore, we evaluated the toxicity using the CellTiter-Glo® viability assay. The toxicity of ICG AGG NPs was compared with that of free ICG following a 4-h incubation with ARPE-19 and MIO-M1 cell lines representing RPE and Müller cells of the retina, respectively (Figure 8A).

**Figure 8.**
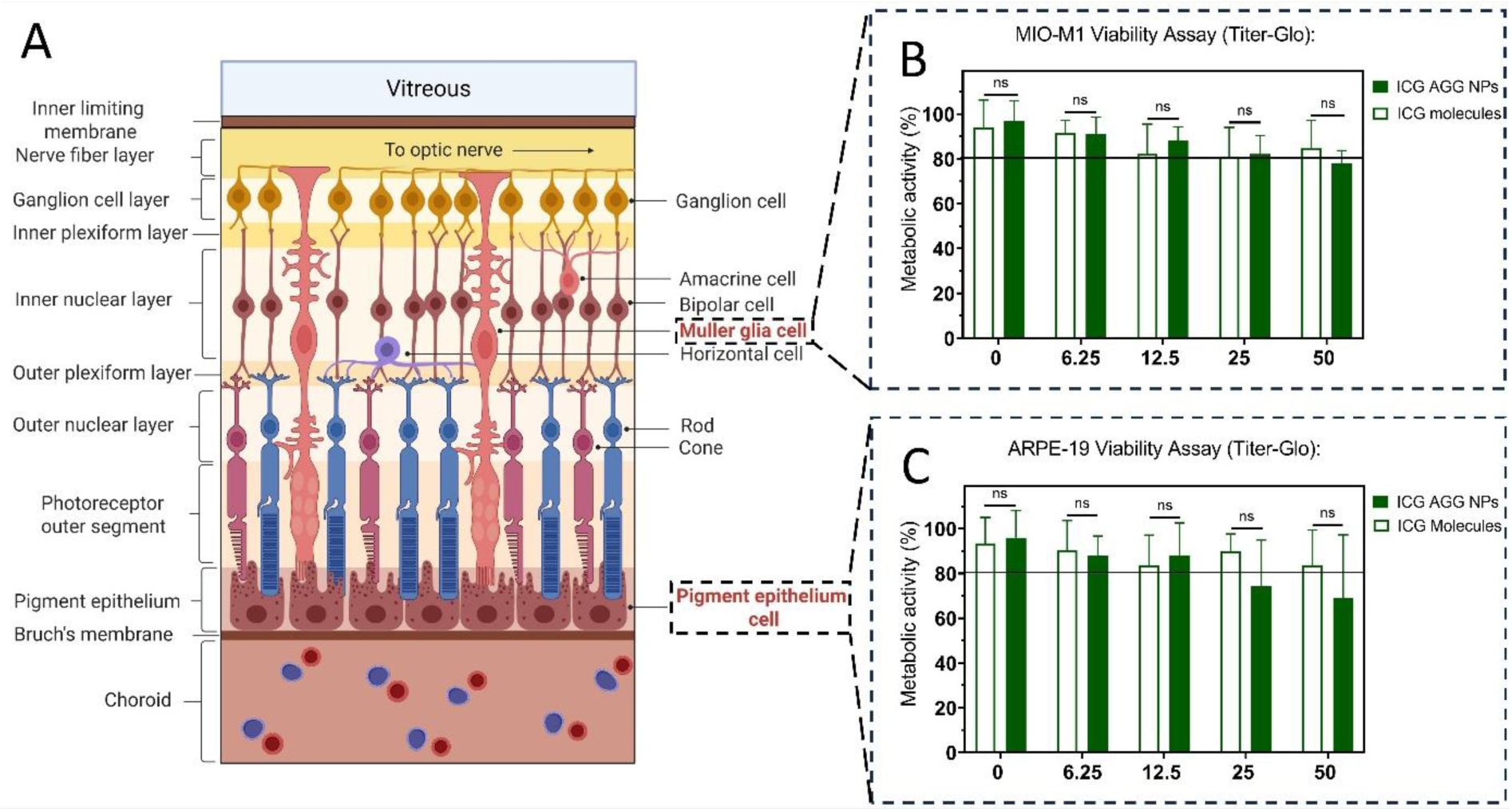
In vitro cell toxicity of ICG AGG NPs. **A)** Schematic representation of the retinal structure, highlighting the retinal pigment epithelium (RPE) and Müller glia cells. The metabolic activity of these two cell types was determined after a 4-h incubation with ICG AGG NPs or free ICG using the CellTiter-Glo® viability assay. **B)** The MIO-M1 cell line derived from Muller glia cells, and **C)** the ARPE-19 cell line, derived from RPE cells, were used for these assessments. The experiment was performed in triplicate (n= 3) for each cell type, with four technical repeats (n= 4) for each concentration.; unpaired t-test P-value: ˃ 0.05 not significant (ns). Image created with BioRender.com.

First, toxicity was assessed on Müller cells (MIO-M1), which expand almost through the entire retina from the ganglion cell layer to the outer nuclear layer. Therefore, they represent one of the first cell types likely to be exposed to our NPs. These experiments revealed that ICG AGG NPs were relatively well tolerated by MIO-M1 cells (Figure 8B), with metabolic activities remaining at approximately 80% or higher across all tested concentrations. No statistically significant differences in toxicity were detected between free ICG and ICG AGG NPs at any of the tested concentrations. The same experiments were performed on ARPE-19 cells, representing RPE cells located in the deepest retinal layer, yielded comparable results (Figure 8C). However, in this case, metabolic activity dropped below 80% for ICG AGG NPs at concentrations of 25 and 50 µg/ml, suggesting that RPE cells may be more sensitive to ICG AGG NPs than Müller cells. This slight toxicity could be related to the HA coating, which may enhance cellular uptake via CD44 receptor interactions.

## Discussion

After intravitreal injection, free ICG can be toxic to the retina [22–24] since it can penetrate through the inner limiting membrane (ILM) due to its small molecular weight and fast diffusion. Moreover, ICG, in its molecular state, cannot form VNBs under typical irradiation conditions with a nanosecond pulsed laser. However, when it accumulates onto collagen fibers, it can generate effective VNBs, leading to photodisruption of vitreous opacities [16]. Therefore, this study aimed at formulating ICG into NPs so as to (i) increase the size of the PS in order to limit its retinal penetration while (ii) keeping the capacity to generate VNBs for effective photodisruption of collagen fibers.

Initially, we intended to employ liposomes as NPs of choice for ICG encapsulation due to their broad use for drug delivery. Liposomes were prepared using the classical thin film rehydration method and purified either by dialysis (Cent -) or by dialysis followed by centrifugation (Cent +). We observed that introducing a centrifugation step and removing aggregates during purification led to a loss of both VNB generation and collagen fiber photodisruption upon 561 nm nanosecond laser excitation. Instead, when liposomes were purified by dialysis alone, these aggregates remained present, leading to successful VNB formation and fiber photodisruption. As shown in Figure 9, these findings highlight the critical role of aggregated ICG in enabling VNB generation and photodisruption, while also demonstrating the limited effectiveness of the liposome formulation alone, despite high ICG encapsulation efficiency by partitioning in the lipid bilayer (Figure 2C,D). We hypothesize that this loss of activity is linked to the absence of ICG aggregation within the liposomal bilayer and core as reported by Yu *et al.* [44].

**Figure 9.**
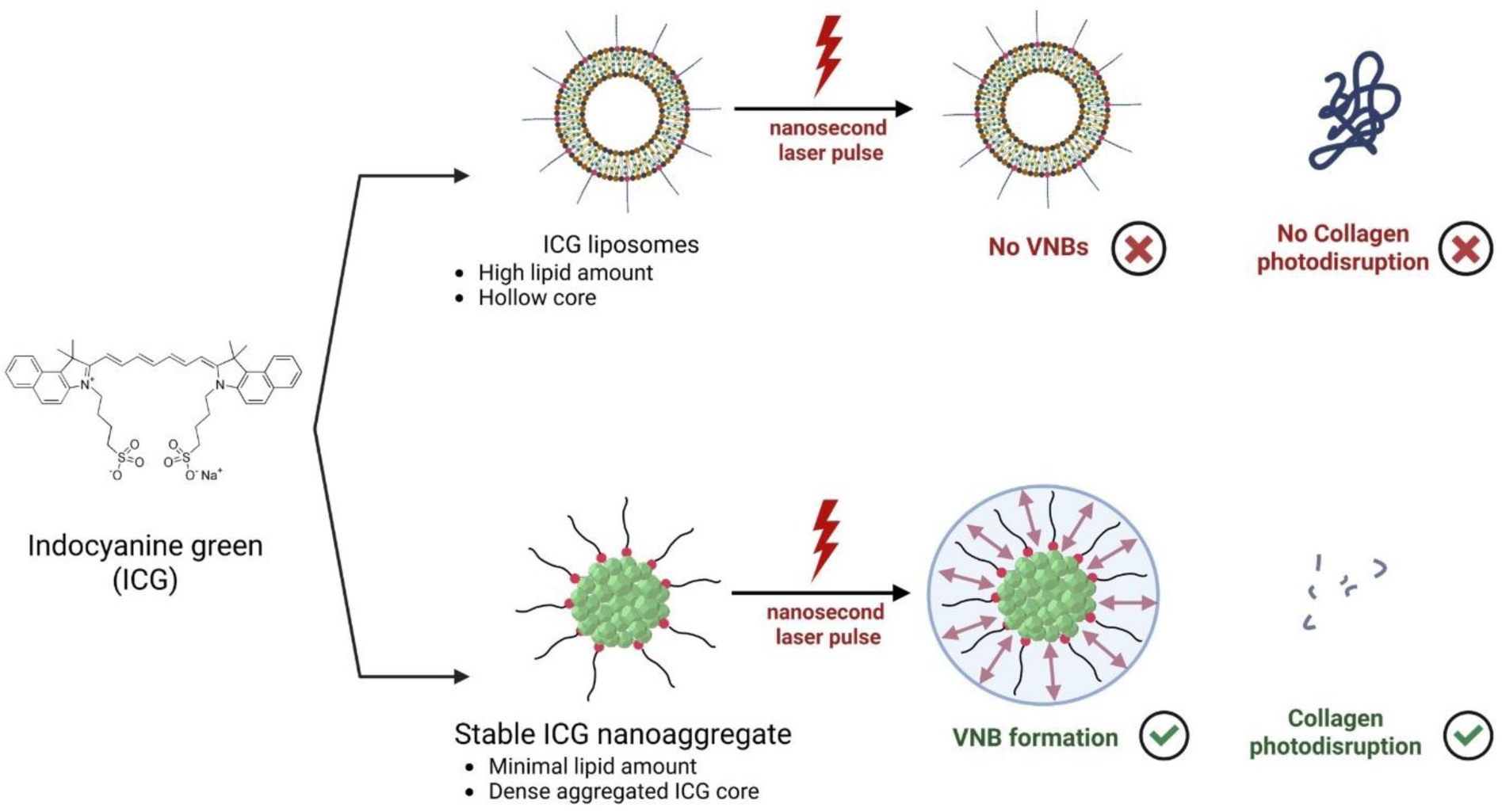
Design and application of ICG NPs for VNB-mediated photodisruption. ICG liposomes were prepared, encapsulating most of the ICG within their lipid bilayer. However, these nanoparticles failed to generate VNBs and could not photodisrupt collagen fibers. In contrast, ICG nanoaggregates with a dense ICG core stabilized by a minimal lipid content efficiently generated VNBs, resulting in effective collagen photodisruption.

Based on the impact of aggregation on VNB generation and photodisruption, we decided to prepare stable nanoscopic ICG aggregates to improve the photodisruption of collagen fibers. Since aggregation is a stochastic and continuous process, controlling and defining the final aggregate size is challenging [45]. Previous studies have addressed this by using lipid- or polymer-based nanocarriers into which ICG can be loaded, either by promoting its aggregation or by encapsulating preformed aggregates. Their focus was to produce NPs containing ordered J-aggregates [40, 46, 47]. In our work, unordered ICG nanoaggregates (ICG AGG NPs), exhibiting aggregation caused quenching (ACQ), were produced (Figure 5C,D), and their application for VNB-mediated photodisruption of vitreous opacities was explored. The strategy to form these NPs was inspired by the emulsification-solvent evaporation technique. The preparation of ICG AGG NPs was achieved by using a small amount of DOPE-HA lipid-polymer conjugate as the stabilizing component. CryoEM analysis confirmed the shape of these NPs to be spherical with a dense core (Figure 5E), in contrast to LIPs, which had hollow cores with ICG accumulation in the lipid bilayer (Figure 2D). The role of DOPE-HA was critical, as it yielded a high number of relatively monodisperse ICG AGG NPs (Figure S3A).

As expected, ICG AGG NPs could successfully generate VNBs and photodisrupt collagen fibers (Figure 9). These NPs were evaluated using two excitation wavelengths of 561 and 800 nm (Figure 6). Remarkably, VNB generation occurred at both wavelengths, with comparable thresholds, requiring fluences below 2 J/cm² and only a few laser pulses (Figure 6A). On average, this corresponds to a total light dose that is 10^3^-10^6^ times lower than typically used in clinical settings for the treatment of vitreous opacities where the number of pulses can reach several hundred to one thousand at significantly higher fluences [16, 48]. Subsequent *in vitro* experiments on collagen fibers confirmed an efficient and successful VNB-mediated photodisruption at both excitation wavelengths (Figure 6B), while no effects were observed when the laser was applied in the absence of ICG AGG NPs (Figure 4C, Figure S4).

In addition to qualitative assessment of fiber photodisruption using dark-field microscopy, we propose a method based on dark field microscopy and light scattering intensity to quantify the photodisruption efficiency of the tested PS. This was achieved by measuring the scattering intensity as a function of the pulse number within a laser-treated area in a “collagen-cloud” model consisting of homogenized collagen aggregates rather than heterogenous thread-like fibers. The collagen cloud was incubated for 24 h with the respective PS before imaging. After treatment with a 561 nm laser, Cent + ICG-HA and ICG-PEG LIPs did not show any photodisruption similar to the control group (without PS). However, free ICG caused a 20% decrease in scattering intensity after 8 pulses. Laser irradiation with ICG AGG NPs showed a significantly better photodisruption capacity as it led to 70% decreasing in scattering intensity after 3 pulses (Figure 7B).

The laser settings that we used for this study are close to the ones used in clinical practice in ophthalmology. First, the pulsed duration (<7ns) is similar to that of the YAG laser used clinically for treating eye floaters. Furthermore, ICG AGG NPs were shown to react to pulsed-laser irradiation at a wavelength of 561 nm, which is close to the 532 nm laser commonly used in clinical settings, e.g. for selective laser trabeculoplasty (SLT) [49]. Also, we could achieve photodisruption with a total applied light dose that is 10^3^-10^6^ times lower than the ones used in the clinics, with a number of pulses not exceeding 30 (while up to hundreds or thousands of pulses are employed for floater destruction with the YAG laser). The ICG concentration at which photodisruption could be observed was much lower than the clinically used 1.25 mg/ml concentration for effective ILM staining [50]. Our *in vitro* toxicity studies also confirmed the safety of these NPs, as no significant toxicity associated with the presence of HA—despite its potential to promote nanoparticle uptake via CD44—was observed (Figure 8). Though these results are promising and demonstrate the use of clinically approved modalities (pulsed-laser, ICG), further studies are needed to assess the efficacy and validate the toxicity of our approach *in vivo*.

## Conclusions

We previously demonstrated the potential of FDA-approved ICG for the photodisruption of vitreous opacities. However, due to its small molecular weight and associated reports of dark- and light-induced retinal toxicity, strategies to limit its retinal diffusion and enhance ocular clearance are needed. Based on previous reports, formulation of ICG into NPs represents a promising approach to mitigate these risks by reducing retinal penetration. However, encapsulating ICG into NPs while keeping their capacity for VNB-generation upon irradiation with a pulsed-laser remains challenging. We found that conventional nanocarriers such as liposomes hinder VNB formation, whereas promoting the aggregation of ICG into dense, solid particles—using only minimal amounts of stabilizing lipid–HA conjugate—yields nanoparticles with strong photomechanical activity. Upon irradiation with a pulsed laser, these ICG AGG NPs achieved a significantly higher photodisruption efficiency compared to molecular ICG. Besides, the encapsulation strategy we employed relies on minimal excipients, which may facilitate industrial scalability.

Interestingly, we also observed that VNB generation is not strictly dependent on the PS’s absorption at the excitation wavelength. ICG AGG NPs generated VNBs with comparable thresholds at both 561 nm (low absorption) and 800 nm (high absorption), suggesting that supramolecular structure and thermal confinement play a more critical role than light absorption.

## Declaration of Generative AI and AI-assisted technologies in the writing process

During the preparation of this work the author(s) used chatGPT4 in order to improve language and readability. After using this tool/service, the author(s) reviewed and edited the content as needed and take(s) full responsibility for the content of the publication.

## Acknowledgements

This work was financially supported by the Phospholipid Research Center (Grant Number SDS-2023-107/1-1). F.S. acknowledges the European Research Council (ERC) for funding received under the European Union’s Horizon Europe research and innovation program (Grant agreement No. 101075873, DYE-LIGHT).

