## Supplementary material for "Lipid-stabilized ICG Nanoaggregates for the Photodisruption of Vitreous Opacities": Videos

### Slide 1
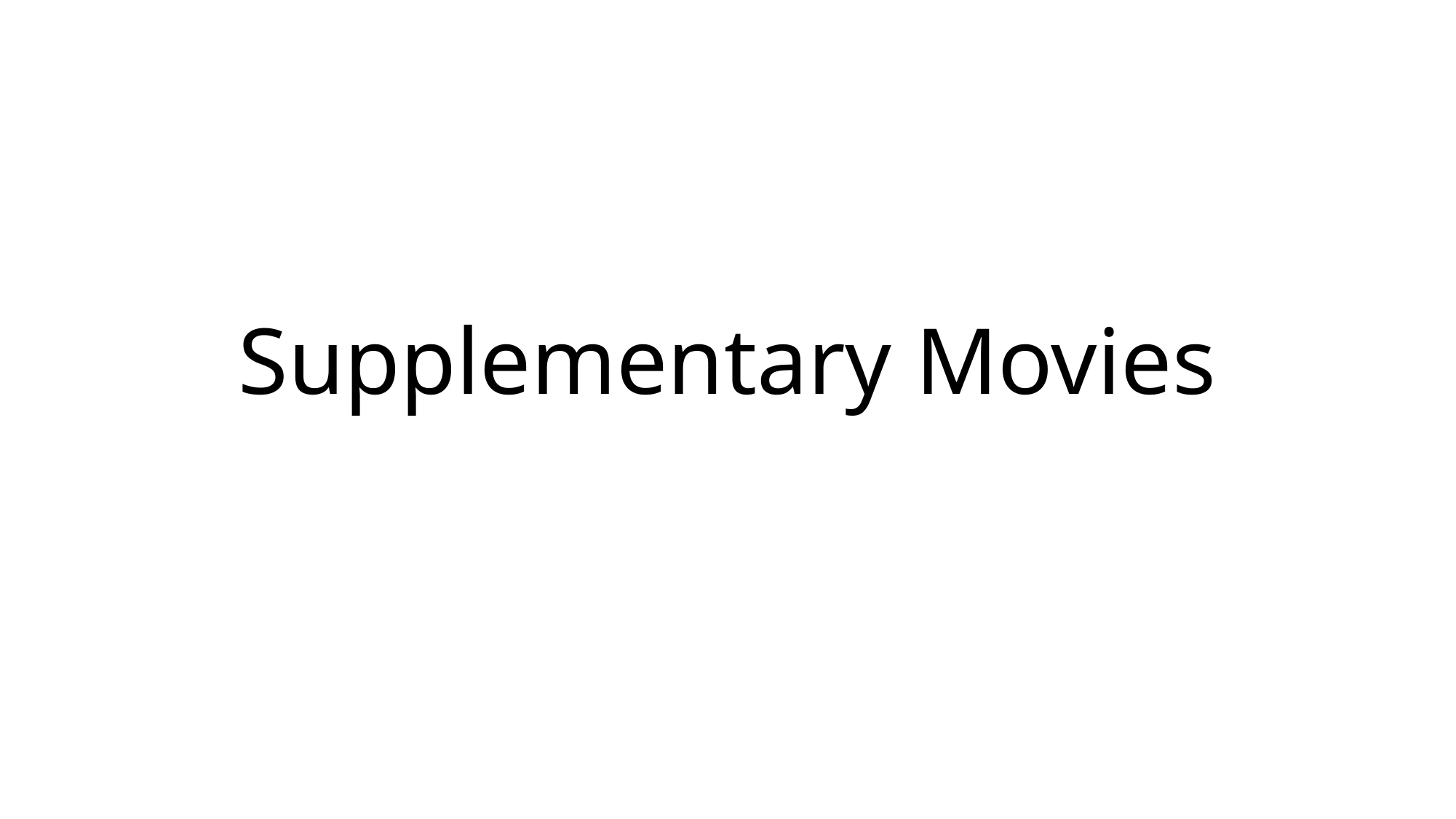

#### Supplementary Movies

### Slide 2
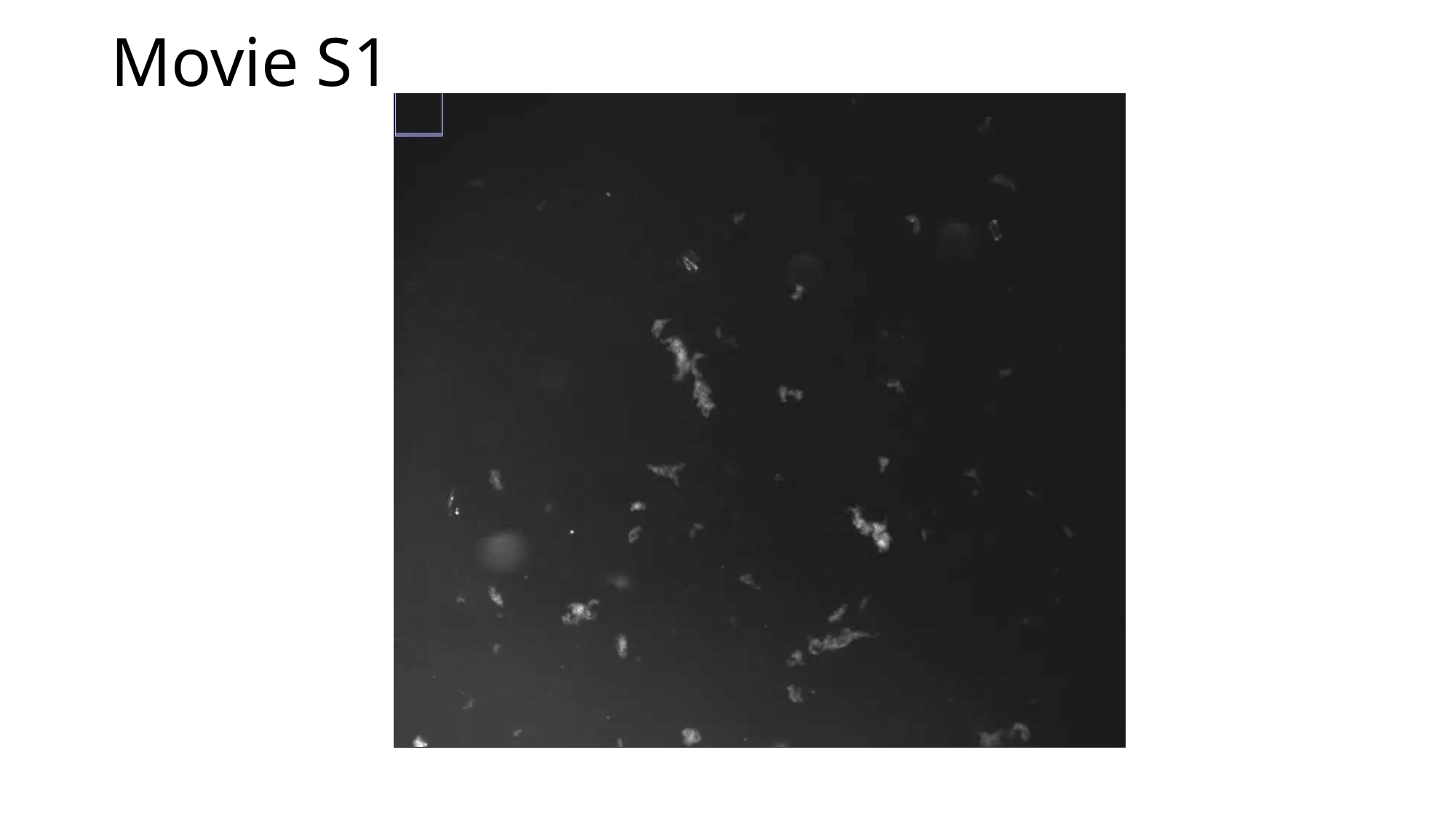

#### Movie S1

### Slide 3
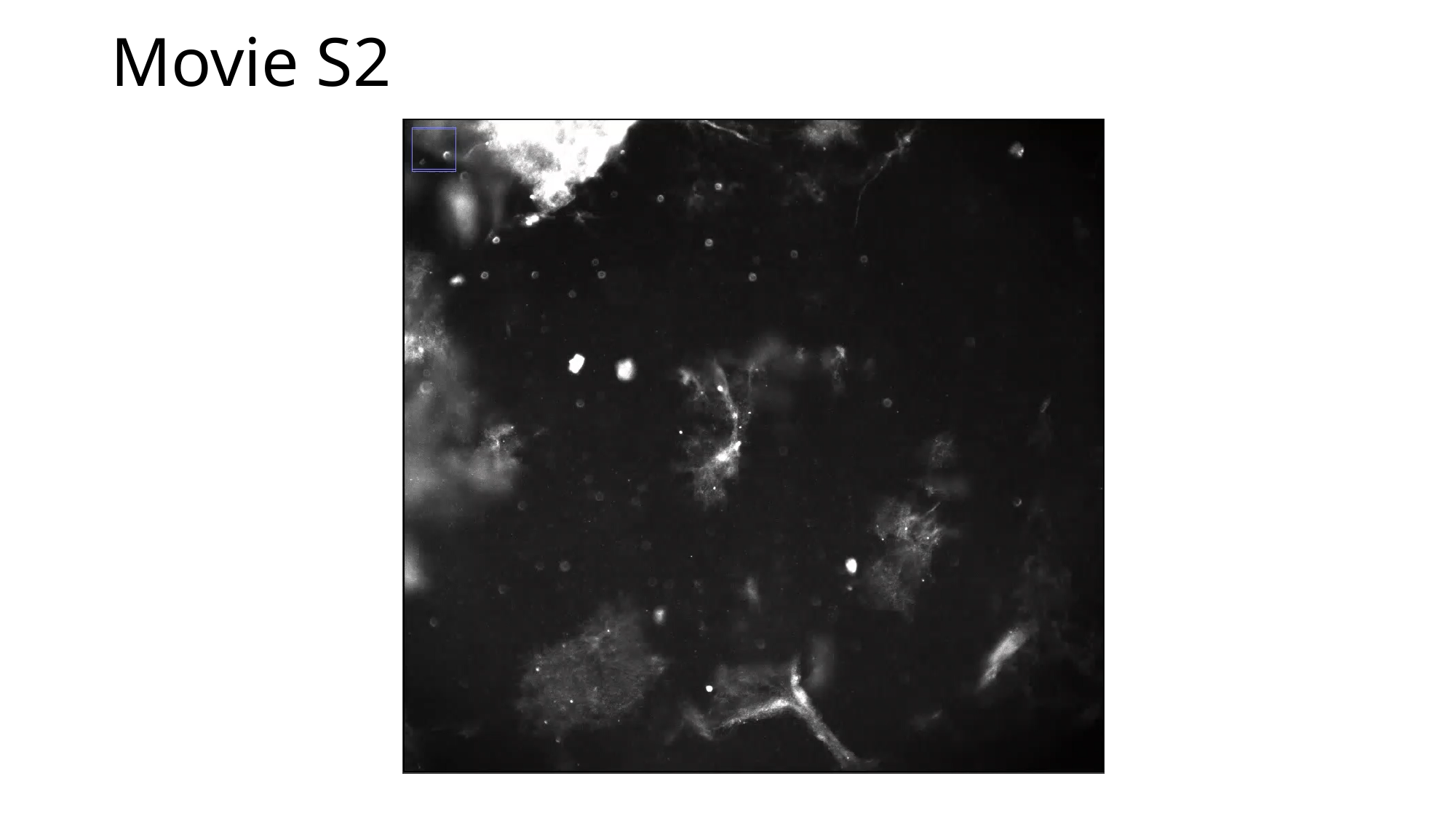

#### Movie S2

### Slide 4
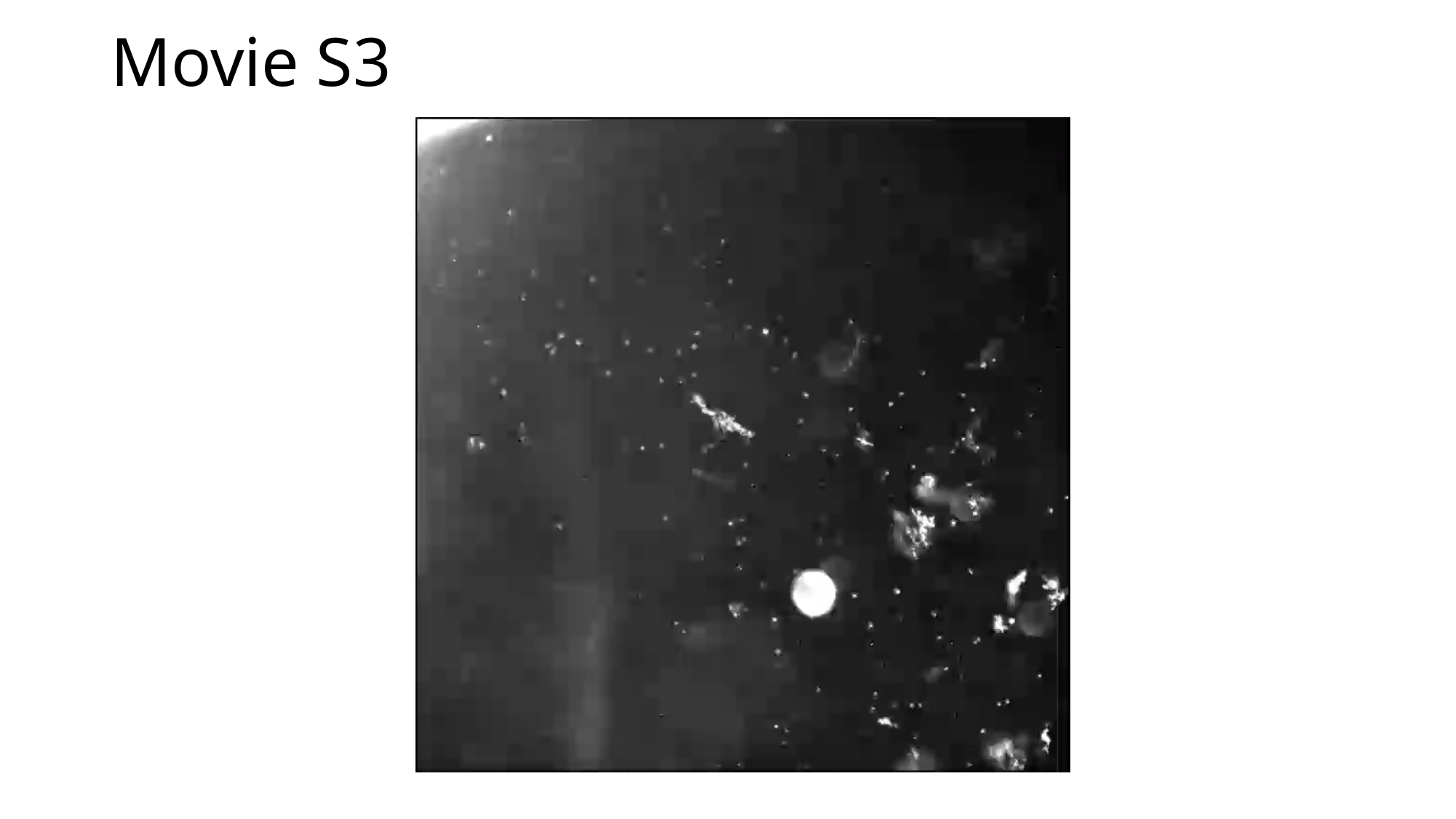

#### Movie S3

### Slide 5
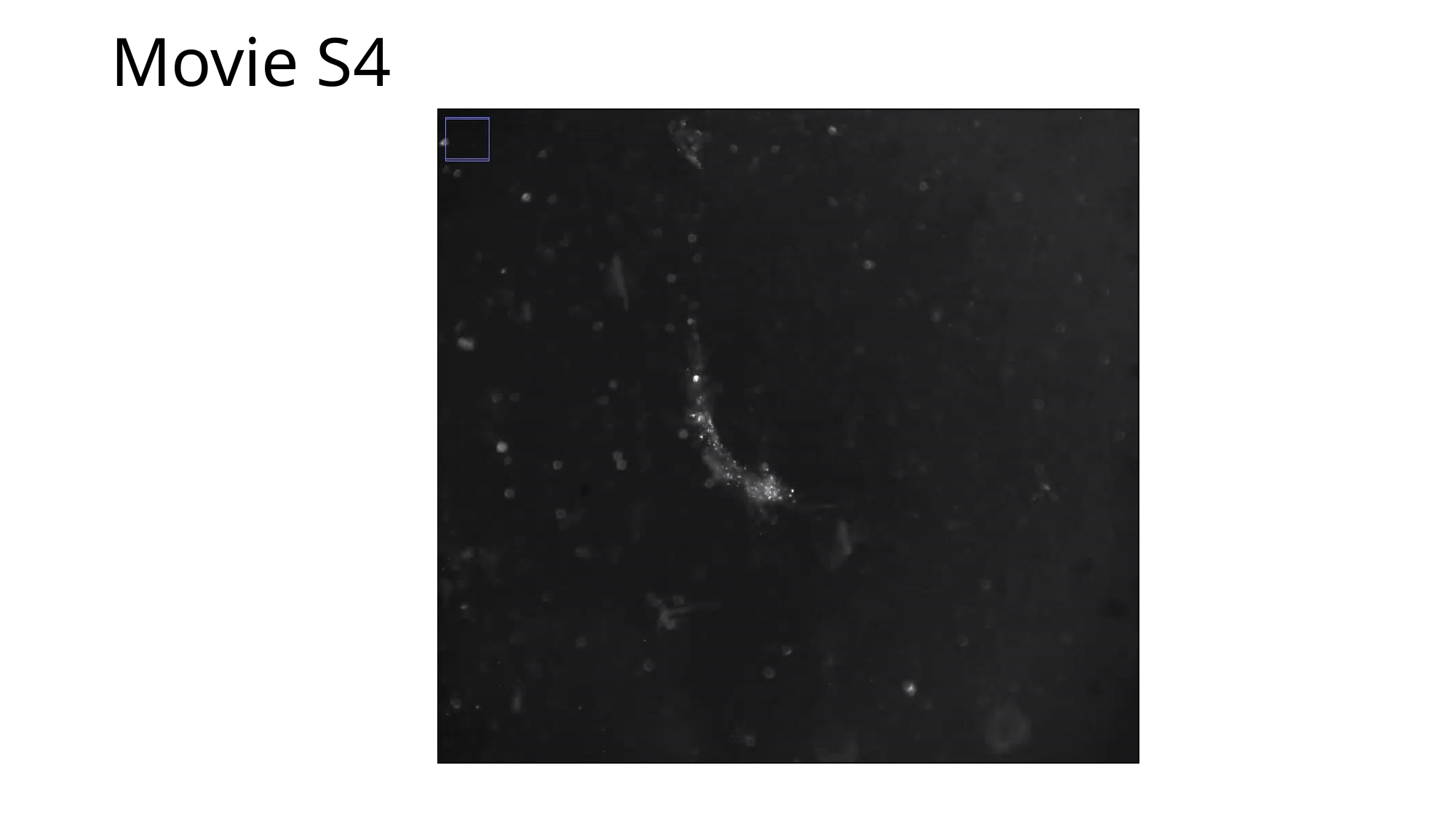

#### Movie S4

### Slide 6
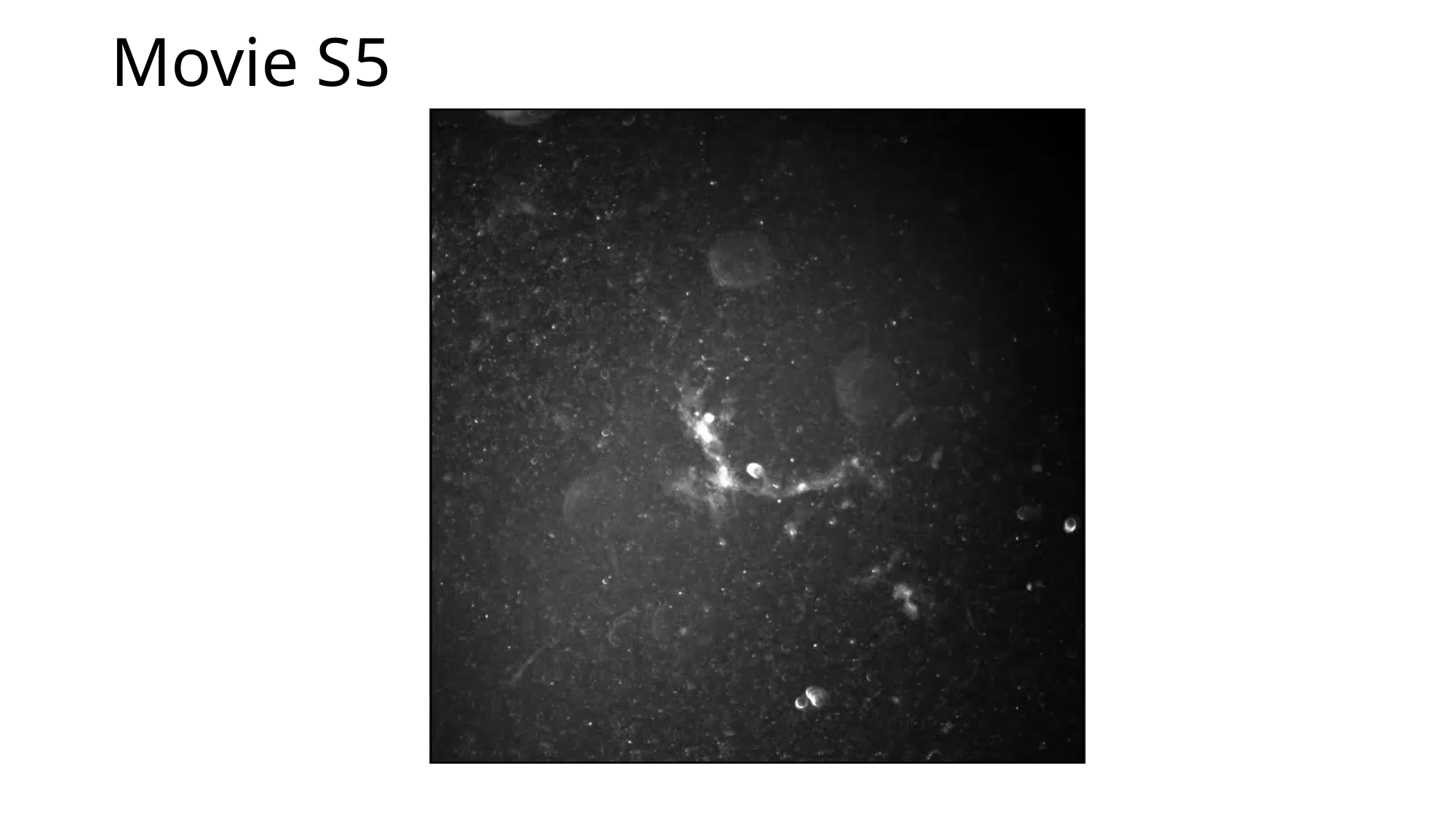

#### Movie S5

### Slide 7
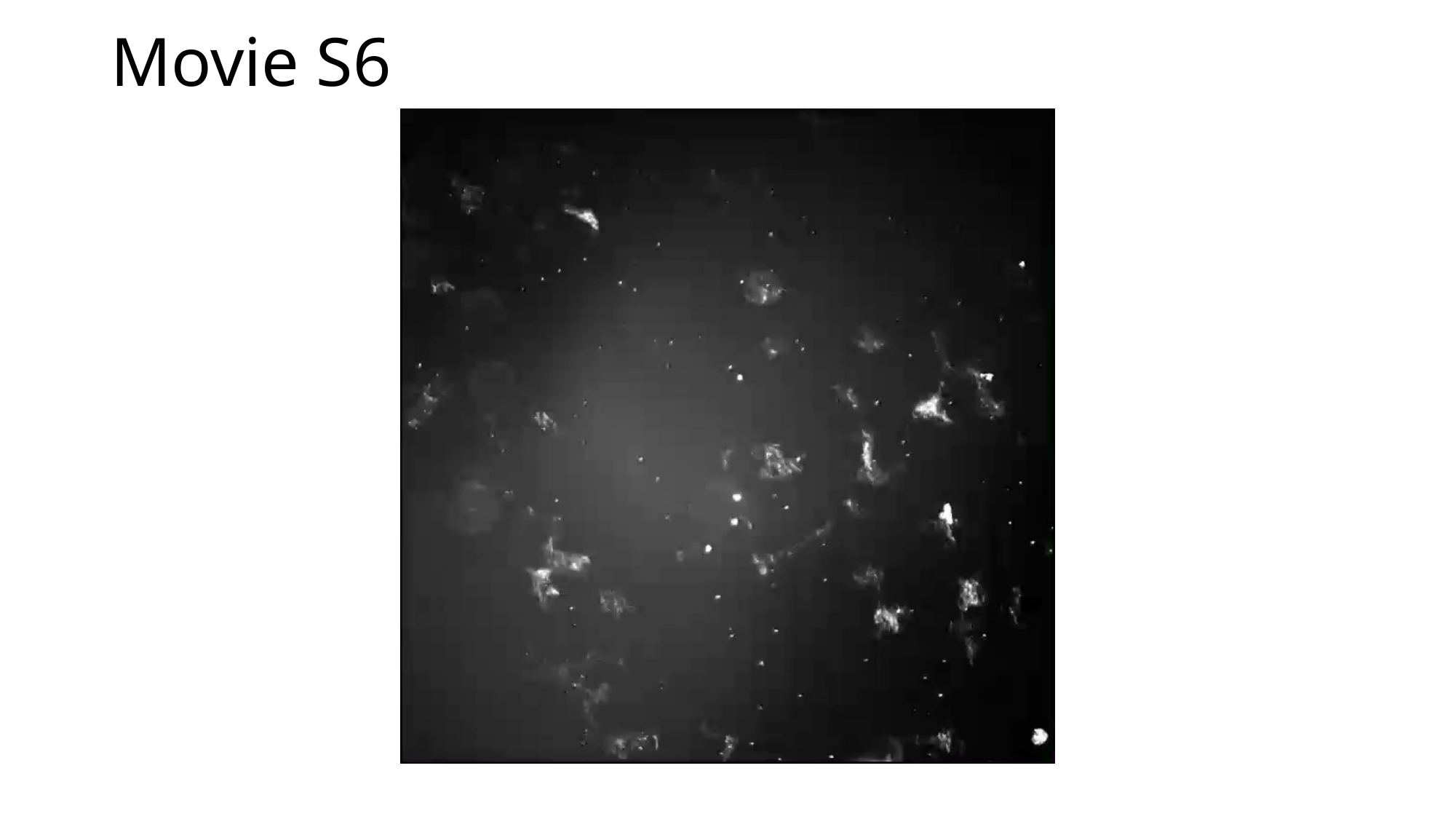

#### Movie S6
