## Supplementary information for "Lipid-stabilized ICG Nanoaggregates for the Photodisruption of Vitreous Opacities"

### *Characterization of DOPE-HA*

The final DOPE-HA product was characterized by measuring the $^{1}H$ NMR spectra (BrukerAV-400 spectrometer) of DOPE-HA compared to unmodified HA, both dissolved in $D_2O$ (Figure S1). In the DOPE-HA spectrum, two new peaks appeared that were absent in the HA spectrum. Peak a, at 0.9 ppm, corresponding to the terminal methyl protons ($–CH_3$) of the DOPE acyl chains and Peak b, at 1.31 ppm (triplet, J = 8.0 Hz), that corresponds to the methylene protons ($–CH_2–$) in the lipid backbone. The degree of substitution (DS) of DOPE on HA was calculated and found to be 9.68%. This was determined similar to Pandolfi *et al.* [1], and by comparing the integrals of the methyl protons of the N-acetyl group in HA (peak c, 2.03 ppm) with those of the terminal methyl groups of DOPE (peak b, triplet, 1.31 ppm).

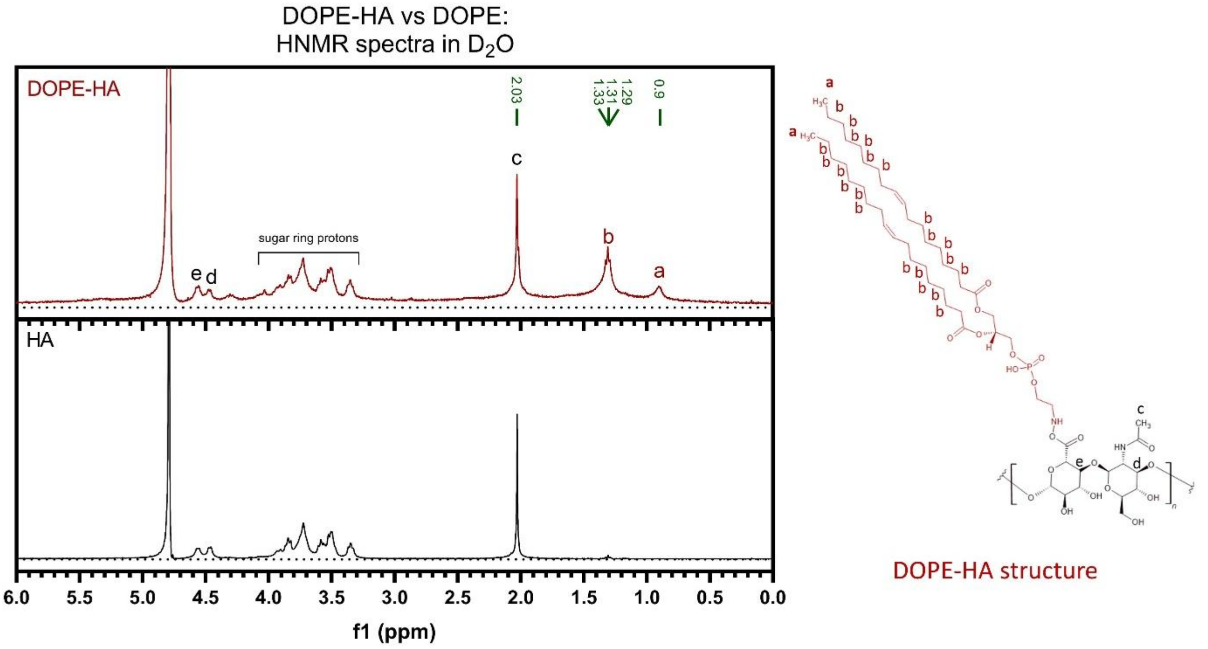


***Figure S1. Characterization of DOPE-HA conjugate by $^{1}$H NMR spectroscopy. Left:*** *$^{1}$H NMR spectra of DOPE-HA (top panel) and pure HA (bottom panel), both recorded in $D_2O$. Based on the DOPE-HA spectrum, the degree of substitution (DS) of the DOPE lipid was calculated to be 9.68%.* ***Left:*** *Chemical structure of DOPE-HA with labeled protons corresponding to the peaks in its $^{1}$H NMR spectrum.*

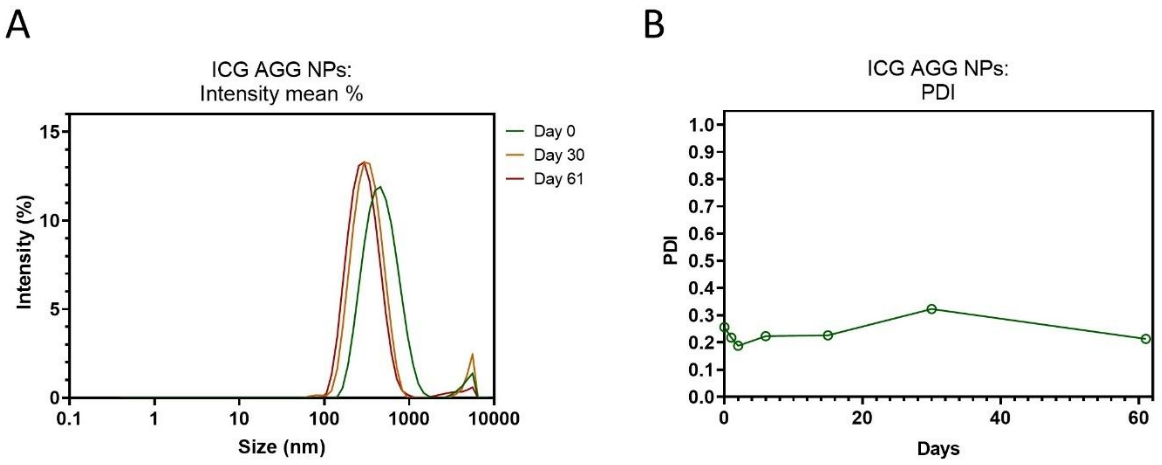


***Figure S2. Stability of ICG AGG NPs evaluated over a two-month period**. Changes in **A)** nanoparticle size, monitored by scattered mean intensity (%,) and **B)** polydispersity (PDI), were evaluated over a two-month storage period in water at 4 °C. All measurements were performed using dynamic light scattering (DLS).*

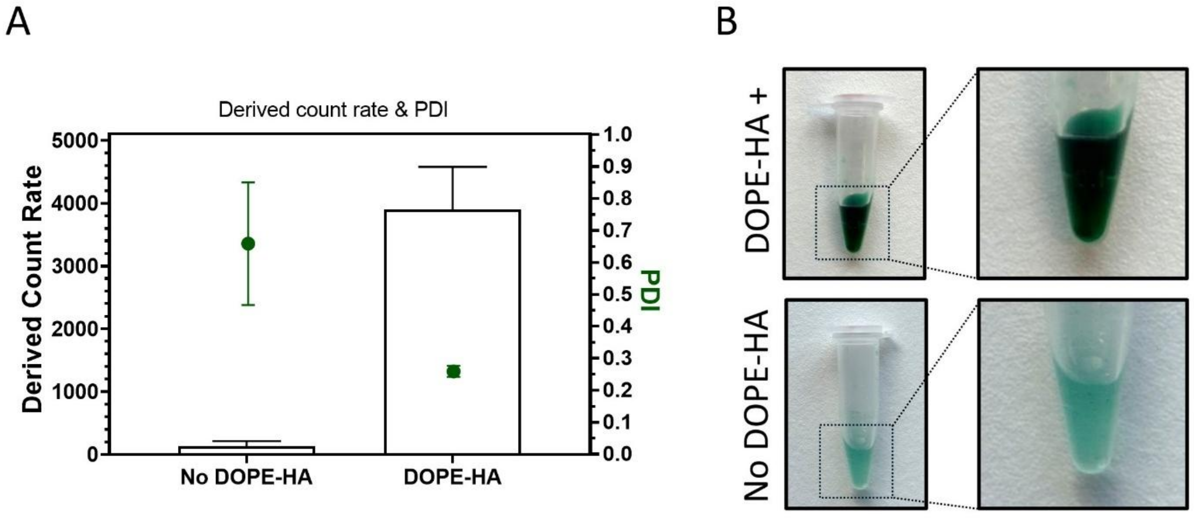


***Figure S3. The effect of DOPE-HA on the polydispersity index (PDI), particle count (derived count rate), and visual appearance of ICG nanoaggregates. A)** Derived count rate and PDI measurements of ICG nanoaggregates prepared with and without DOPE-HA. **B)** Visual inspection of the samples.*

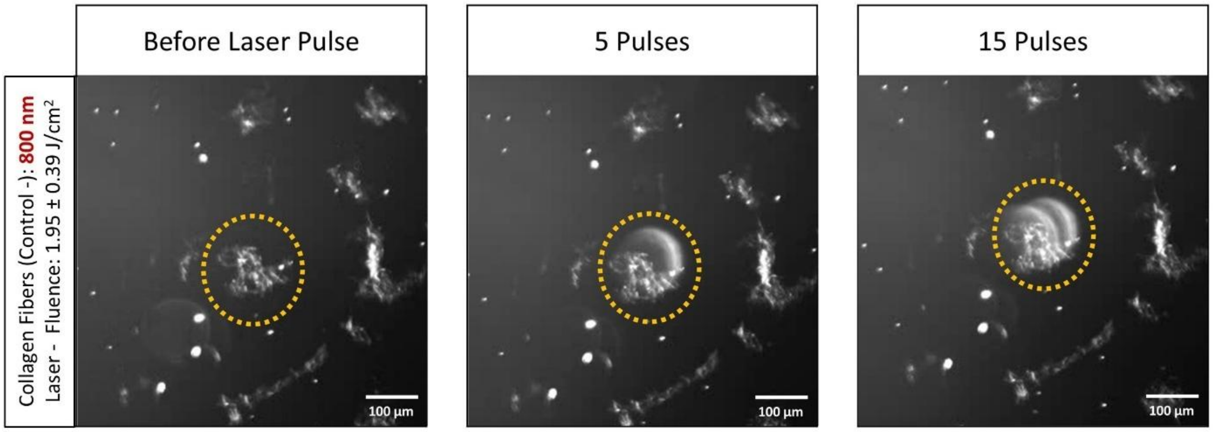


***Figure S4. Representative dark-field image of an artificial collagen fiber before and after 5 and 15 pulses with the nanosecond laser (< 7ns; 800 nm; 1.95 J/cm²)****. Scale bar 100 µm.*

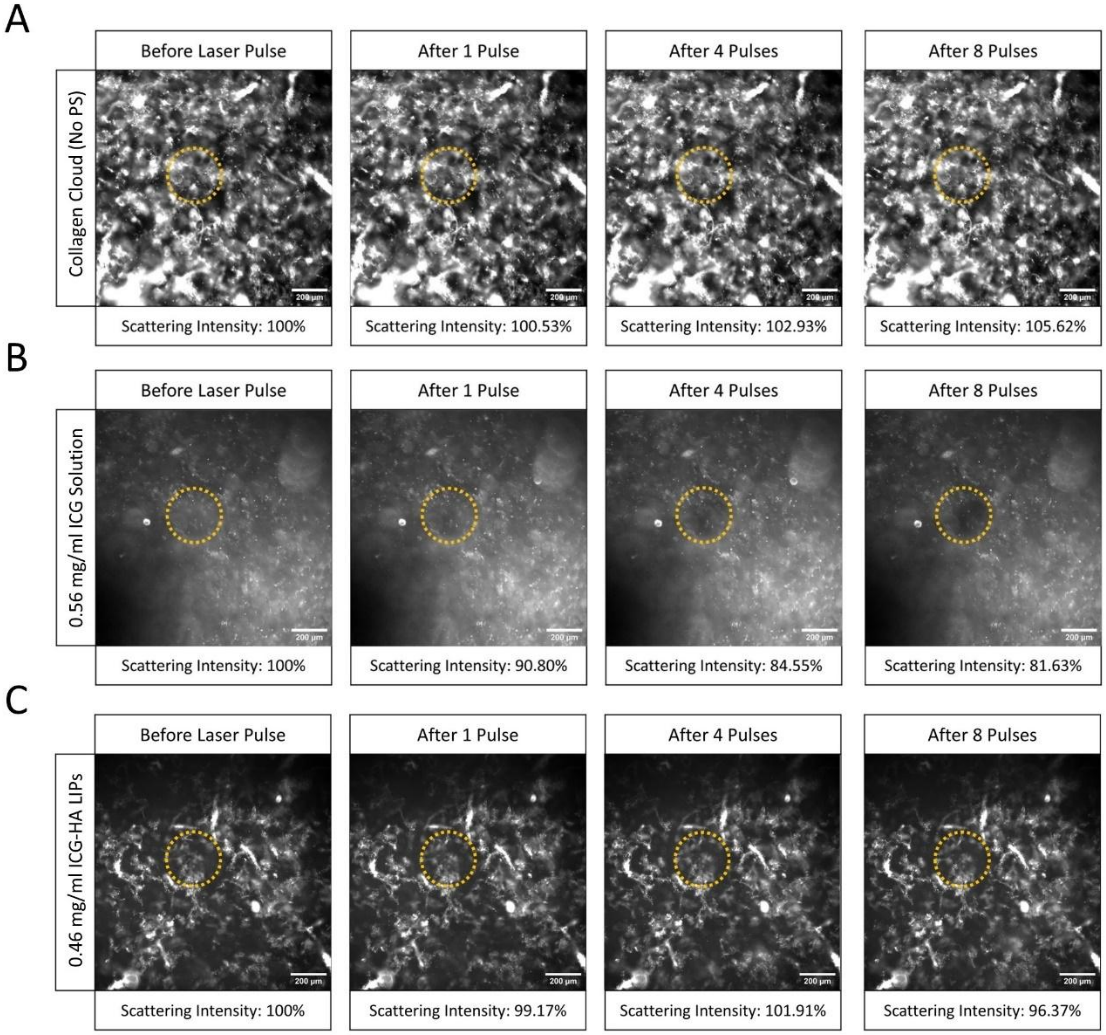


***Figure S5. Representative dark-field microscopy images used for the photodisruption quantification. A)*** *the collagen cloud in the absence of any PS (no PS),* ***B)*** *free ICG, and* ***C****) ICG-HA LIPs. Scale bar 200 µm.*

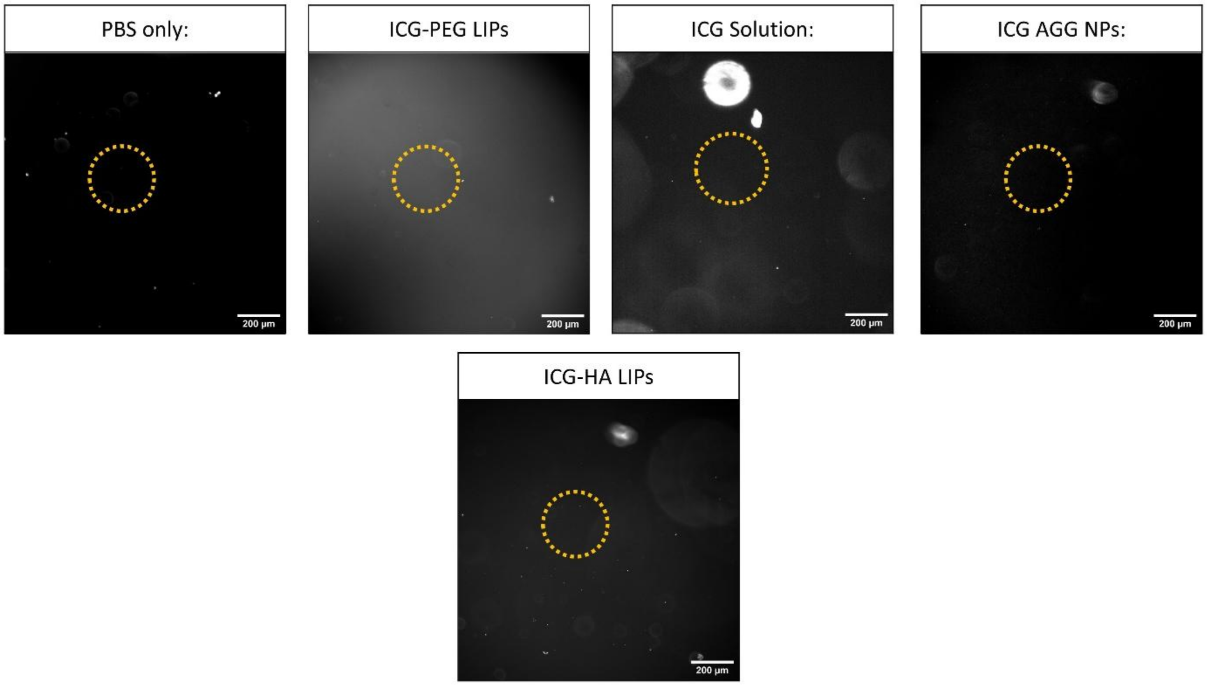


***Figure S6. Dark-field images of photosensitizer formulations without collagen, used as reference for baseline scattering intensity.*** *These images served as the baseline* ($II_0$) *for accurately calculating collagen cloud scattering intensity (Figure 7B). Scale bar: 200 µm.*
